# Enhancement of Chikungunya virus genome replication in mammalian cells at a sub-physiological temperature

**DOI:** 10.1101/2021.09.29.462334

**Authors:** Jinchao Guo, Aaron E. Lin, Chetan Aditya, Alexander Ploss, Mark Harris

## Abstract

Chikungunya virus (CHIKV) is an alphavirus transmitted by *Aedes* mosquitoes, causing fever, rash and arthralgia in mammals. The function of the CHIKV non-structural protein 3 (nsP3) remains enigmatic. Building on previous studies (Gao *et al*, 2019) [1], we generated a panel of mutants in a conserved and surface-exposed cluster in the nsP3 alphavirus unique domain (AUD) and tested their replication using a sub-genomic replicon (SGR). Three of these SGR mutants replicated well in mosquito cells but poorly in mammalian cells. We observed that this difference was due to culture temperature (mosquito cells: 28°C; mammalian cells: 37°C), as the mutants exhibited no replication defect in mammalian cells grown at a sub-physiological temperature (28°C). Similar phenotypes were observed for infectious CHIKV and the closely related O’Nyong Nyong virus. Intriguingly, the wildtype SGR replicated more efficiently in mammalian cells at 28°C compared to 37°C. To explore the mechanism behind this difference, we focused on two known antiviral pathways: interferon-stimulated genes (ISGs) and stress granules (SGs). SGR replication was concomitant with increased expression of ISGs at 37°C, but this response was impaired at 28°C. We also observed enhanced recruitment of the SG component G3BP1 into cytoplasmic sites of viral genome replication at 28°C. These findings may have real-world implications as when a mosquito bites a mammal, the virus first infects cells in peripheral tissues which are often at sub-physiological temperatures. We propose that alphaviruses such as CHIKV have evolved mechanisms to both promote viral genome replication and concomitantly limit antiviral responses in these cells.

**Significance statement:** Chikungunya virus (CHIKV) is a re-emerging arbovirus that is transmitted by *Aedes* mosquitos and poses epidemic threats. Arboviruses must be able to grow efficiently in both the mosquito vector and the mammalian host, which have different body temperatures. Following a mosquito bite, the first cells infected are typically in the skin and are at sub-physiological temperatures, ie lower than the internal body temperature (37°C). Here we show that CHIKV grows more efficiently in mammalian cells cultured at 28°C, compared to 37°C. We investigated the mechanism behind this increase in growth efficiency and demonstrated that at the lower temperature the production of antiviral genes was impaired. Additionally, proteins that respond to cellular stress and are needed for virus replication were more abundantly recruited to sites in the cell where the virus was replicating. We propose that CHIKV has evolved mechanisms to promote its replication in mammalian cells at sub-physiological temperatures to facilitate infection of mammals via a mosquito bite.

## Introduction

Chikungunya virus (CHIKV) belongs to the *alphavirus* genus in the *Togaviridae* family [2]. It is an arbovirus that is transmitted by mosquitoes and causes fever, rash and arthralgia with infrequent long-term morbidity and, in rare cases, mortality in mammals [3]. CHIKV was first isolated in Tanzania in 1952-1953 [4, 5], and has subsequently spread across the world, including Africa, Europe, Asia and America [6, 7]. CHIKV is mainly transmitted by the mosquito species *Aedes (Ae.) aegypti* and *Ae. albopictus*, and the latter has become the main vector for virus spread in Asia [8]. Both field and laboratory experiments demonstrated that these two mosquito species only survive between lower and upper temperature thresholds [9]. Models and human cases showed that transmission occurs between 18°C and 34°C with maximal transmission occurring in a range from 26-29°C [10].

CHIKV has a positive-sense, single-stranded RNA genome, 11.5 kb in length, which is both capped and polyadenylated (Fig. S1A). It contains two open reading frames (ORFs). The first ORF encodes the non-structural proteins (nsP1-nsP4) and is translated directly from full-length genomic RNA. ORF2 encodes the structural proteins (capsid, E1-E3 and 6K) and is translated from a sub-genomic RNA. The non-structural proteins are required for the synthesis of positive and negative sense full-length genomic RNA as well as the sub-genomic RNA. Biochemical analysis indicates specific functions of the non-structural proteins: nsP1 has methyl- and guanyl-transferase activities, nsP2 exhibits helicase and protease activities and nsP4 is the RNA-dependent RNA polymerase [11].

The function of nsP3 remains poorly defined [12], despite being essential for genome replication. CHIKV nsP3 consists of three domains: a macro-domain at the N-terminus, a central alphavirus unique domain (AUD) and a hypervariable domain (HVD) at the C-terminus (Fig. S1A). The macro-domain exhibits RNA binding, ADP-ribose binding and ADP-ribosylhydrolase activities [13–15]. The HVD is highly phosphorylated and interacts with a number of host proteins [16]. Despite high levels of conservation across alphaviruses, AUD function remains elusive. In this context, we showed that the AUD binds to viral RNA and plays a role in virus assembly by stimulating transcription of the sub-genomic RNA and concomitant translation of ORF2 [1]. In addition, recent cryo-EM analysis revealed that nsP3 assembles into tubular structures, driven by a helical arrangement of the AUD [17]. AUD mutations that disrupted these assemblies also impaired virus replication.

nsP3 also plays multiple roles in rewiring host cell biology, especially altering stress granules (SGs) [18]. SGs are cytoplasmic, non-membranous aggregates formed by eukaryotic cells in response to cell stress and translation inhibition. Typical SGs contain silent mRNAs, translation factors, small ribosomal subunits, and various mRNA-binding proteins. Despite varied compositions resulting from different stresses [19], the RasGAP SH3-domain binding proteins (G3BP1/2) are key components required for formation of all SGs [20–22]. However, alphavirus infection blocks SG formation: the CHIKV nsP3 hypervariable domain interacts with G3BP1/2 and sequesters them into cytoplasmic foci which are the sites of genome replication [23, 24] [18]. Recent work also demonstrated that the CHIKV nsP3 macro-domain can alter the composition of SGs or disassemble them entirely [25]. Therefore, CHIKV nsP3 plays a key role in both blocking SG formation and altering their composition in order to promote genome replication.

In addition to altering SGs, CHIKV must also limit the interferon (IFN)-mediated innate immunity, which provides the first line against invading pathogens. As with many RNA viruses, CHIKV genome replication produces double-stranded RNA which is sensed by infected cells, which in turn produce and secrete type I IFNs to induce an antiviral state in neighboring cells [26]. Cells that sense type I IFNs initiate the JAK-STAT signaling pathway and the downstream production of IFN-stimulated genes (ISGs) [27]. This leads to antiviral states which are effective in restricting replication of both DNA and RNA viruses [28]. A number of ISGs have been shown to suppress infection by the model alphavirus Sindbis virus [29]. Additionally, it has been shown that in a mouse model production of type I IFN was impaired at sub-physiological temperatures, and thus CHIKV replication and arthritis was more pronounced [30, 31].

In this study we sought to investigate the function of the nsP3 AUD in CHIKV replication in cells from both the vertebrate host and the mosquito vector. Three AUD mutants (W220A, D249A and Y324A) replicated poorly in mammalian cells but efficiently in mosquito cells. However, culture temperature confounded this result, since mammalian cells are typically cultured at 37°C while mosquito cells are cultured at 28°C. We demonstrate that a wildtype CHIKV sub-genomic replicon (SGR) replicated more efficiently in mammalian cells at a sub-physiological temperature, suggesting that the original AUD mutant phenotype was temperature-dependent rather than species-specific. To explore the mechanism behind differential replication at 28°C or 37°C, we focused on two known antiviral pathways: ISGs and SGs. Transcriptomic analysis revealed that ISG expression was impaired at the lower temperature. Furthermore, analysis of SGs revealed that G3BP1 was more efficiently recruited to cytoplasmic replication sites at the sub-physiological temperature. Taken together these observations may explain, at least in part, why replication of CHIKV SGR in mammalian cells is enhanced at a lower temperature.

## Results

### Construction and analysis of AUD mutants

In this study we set out to build on our previous work which had identified a role for two residues (P247 and V248) in the function of the CHIKV nsP3 AUD [1]. Alanine substitution of both residues disrupted the binding of the AUD to the sub-genomic promoter, reducing transcription of the sub-genomic RNA and translation of structural proteins, with concomitant effects on virus assembly. Interestingly, the P247A/V248A mutation also disrupted the helical arrangement of nsP3 observed by cryo-EM [17]. To further investigate the phenotype that we observed, we first generated a series of mutations in surface-exposed and conserved residues proximal to P247 and V248. To identify such residues, AUD amino acid sequences from both Old and New World alphaviruses were aligned (Fig. S1B). The nsP2/nsP3 protein structure of Sindbis virus (SINV) has been determined [32], and the AUD sequences between SINV and CHIKV share high similarity (118 out of 243 residues are identical). A panel of conserved and surface exposed residues in the AUD were therefore selected for mutation based on the SINV nsP2/nsP3 protein structure [32] (Figs. S1C). Seven residues were chosen for further study as they were either identical or similar between the two viruses. The C-terminal residue of nsP3 (Y324 in nsP3 amino acid numbering) was selected as a control as it is distal to P247/V248 and highly conserved among alphaviruses (Fig. S1B).

These mutations were introduced into a CHIKV dual luciferase sub-genomic replicon (CHIKV-SGR-D-Luc) (Fig. S1A), which was derived from the ECSA strain LR2006 OPY1 isolate [33, 34]. In this construct, renilla luciferase (R-Luc) is fused within the nsP3 hypervariable domain in ORF1, and firefly luciferase (FF-Luc) replaces the structural proteins in ORF2. The activity of R-Luc is thus a marker of ORF1 translation from the full-length RNA: at early time points R-Luc levels reflect transfection efficiency and input RNA translation, at later times R-Luc is also dependent upon genome replication. As ORF2 is translated from a sub-genomic RNA (sgRNA) which is templated from the negative strand replicative intermediate, FF-Luc expression is dependent upon genome replication but will also reflect translation from the sgRNA. A polymerase-inactive mutant (GDD>GAA in nsP4) was used as a negative control.

### AUD mutants exhibit different replication phenotypes in mammalian and mosquito cells

CHIKV mainly induces arthritis and joint pain [3] and infects muscle cells. We therefore initially chose to investigate the effects of the AUD mutants on CHIKV genome replication in C2C12 cells, an immortalized mouse myoblast cell line widely used in the study of CHIKV [35]. As expected, wildtype (WT) CHIKV-SGR-D-Luc showed a very high level of replication in C2C12 cells, with FF-Luc levels peaking at 12 h post transfection (h.p.t) and increasing over 1000-fold between 4-12 h.p.t. (Fig S2A). AUD mutants T218Q, M219Q, P221A, C246A and A251Q exhibited similar levels of replication as compared to WT (Fig S2A), however, W220A, D249A and Y324A showed significantly lower levels of replication compared to WT (Fig S2A).

As CHIKV is transmitted by mosquitoes, we then evaluated the replication capacities of the AUD mutants in C6/36 and U4.4 cells, both derived from *Ae. albopictus*. C6/36 cells harbor a frameshift mutation in the Dicer-2 (Dcr2) gene, which leads to the translation of a truncated Dcr2 protein and a defect in the RNAi response [36]. Consistent with this, WT CHIKV-SGR-D-Luc showed robust replication in both cell lines, with higher levels of replication in C6/36 than U4.4 cells (Fig. S2B/C). Interestingly, unlike the replication defects observed in C2C12 cells, all AUD mutants showed similar levels of replication compared to WT in both C6/36 and U4.4 cells (Fig. S2B/C). In C6/36 cells levels of FF-luc continued to increase up to 48 h.p.t. whereas they peaked at 24 h.p.t. in U4.4 cells.

The striking phenotypic differences between mammalian and mosquito cell lines for the three AUD mutants (W220A, D249A and Y324A) led us to investigate further. A simple explanation might be that these mutants had reverted to WT in mosquito cells, as we had previously observed this for the AUD mutant R243A/K245A [1]. This mutant replicated poorly in mammalian cells but efficiently in mosquito cells, and this was explained by the fact that R243A/K245A had reverted to WT in mosquito cells allowing efficient replication. To test this hypothesis, total RNA from U4.4 and C6/36 cells at 72 h.p.t was extracted, the nsP3 coding sequence was amplified by RT-PCR and Sanger sequenced. Reassuringly, none of the AUD mutants had reverted to WT, and all three mutants retained the same sequence as the input RNA (Fig. S2D). The results demonstrated that the ability of these three AUD mutants to replicate in mosquito cells was not due to reversion. We conclude that the three residues W220, D249 and Y324 within the CHIKV nsP3 AUD play a role in genome replication in mammalian cells but not in mosquito cells.

### The AUD mutants exhibit a temperature dependent phenotype

We reasoned that a simple explanation for the different phenotypes in mammalian and mosquito cells might be the temperature difference, as mammalian cells are cultured at 37°C whereas mosquito cells are cultured at 28°C [37]. To test this, the replication of WT and mutant CHIKV-SGR-D-Luc were analysed in C2C12 cells incubated at 28°C. Intriguingly, none of the three mutants exhibited any phenotype at this lower temperature, and FF-luc (Fig. 1B) levels were indistinguishable from WT at 28°C. In addition, we noted that at 37°C WT CHIKV-SGR-D-Luc FF-luc levels reached a plateau at 12 h.p.t. (Fig 1A), but in contrast, at 28°C FF-luc levels continued to increase (approximately 100-fold) from 12 to 24 h.p.t. (Fig 1B). As FF-luc levels were similar at both temperatures up to 12 h.p.t. this suggested that initial rates of replication and/or translation were temperature-independent, however after 12 h.p.t. these rates were limited at 37°C, but this limitation was relieved at the lower temperature. Translation from the CHIKV SGR at the two temperatures in C2C12 cells was quantified by western blotting for nsP1 expression (Fig. 1C). Reassuringly, none of the mutants exhibited detectable nsP1 at 37°C whereas a low level of nsP1 was detectable for the WT at this temperature. In contrast, WT and all mutants exhibited high and equivalent levels of nsP1 at 28°C, consistent with the elevated levels of replication and translation at this temperature.

**Figure 1.**
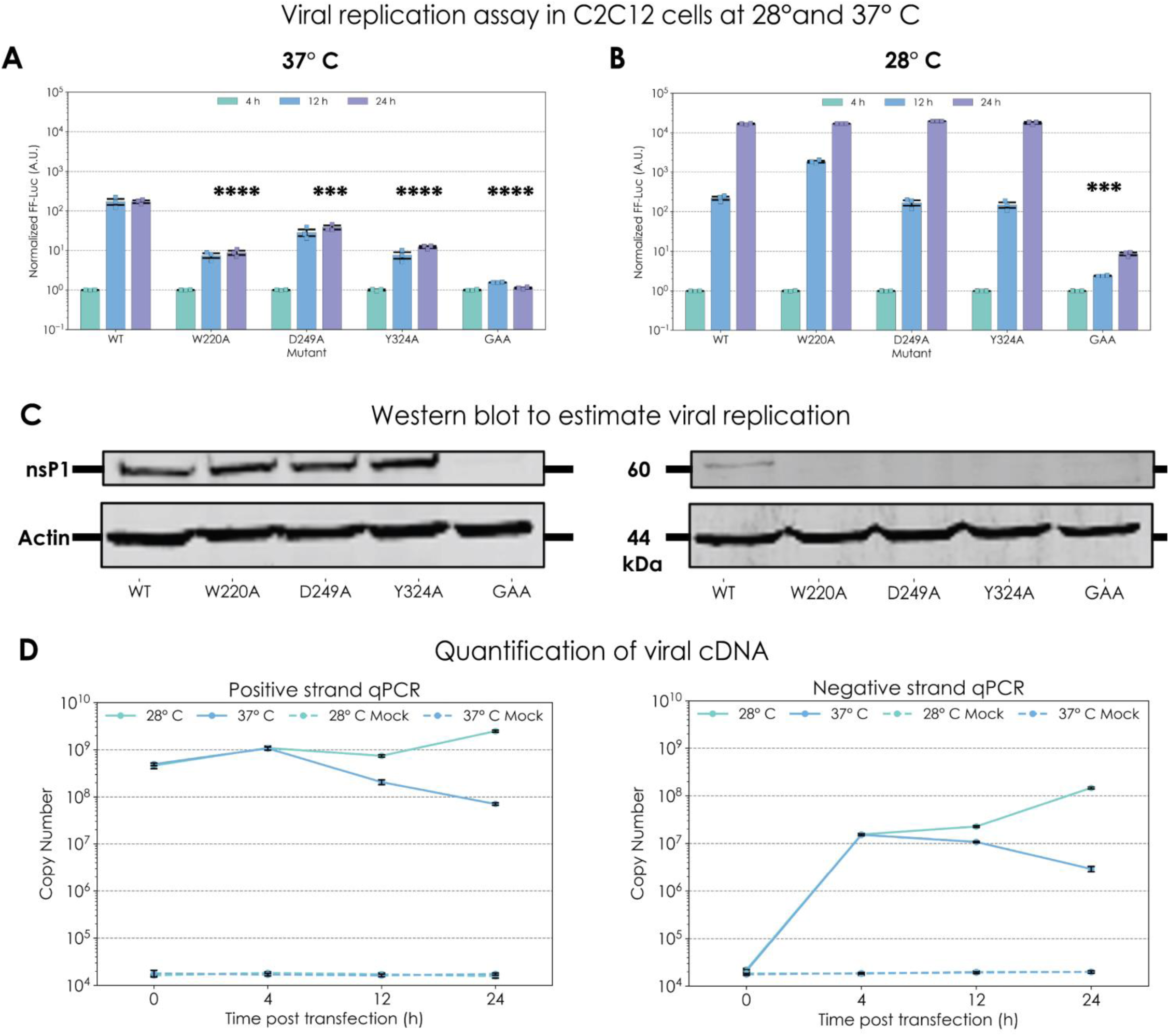
Replication and translation of CHIKV AUD mutants in C2C12 cells at 28°C or 37°C. C2C12 cells were transfected with WT and mutant CHIKV-SGR-D-Luc RNAs and cultured at 37°C (A) or 28°C (B). Cells were harvested at indicated time points. FF-Luc activities of WT and mutants were measured and normalized to their respective 4 h values, which thus represent fold-change compared to the 4 h values. (**C**) Protein expression of nsP1 for the WT and mutant CHIKV-SGR-D-Luc at 28°C or 37°C in C2C12 cells. Cells were lysed at 24 h.p.t., 10 µg of each sample was loaded and analysed by western blotting. Actin was also detected by western blot as a loading control. (**D**) Quantification of CHIKV positive and negative strand RNA from C2C12 cells transfected with CHIKV-SGR-D-luc. Transfected C2C12 cells from (**A**) were lysed with TRIzol solution, and total RNA was purified by phenol-chloroform extraction. cDNAs of CHIKV positive or negative strand RNA were synthesized as described before [64].

To provide further evidence for the impaired replication at 37°C compared to 28°C, we quantified both positive and negative strand CHIKV RNA over a period of 24 h.p.t. in C2C12 cells (Fig. 1D). Although the high level of transfected input RNA hampered quantification of genome replication at early time points (e.g. 4 h.p.t.) a clear distinction in RNA levels could be seen at later time points. Specifically, the levels of both positive and negative strand RNA declined at 37°C between 4-24 h.p.t. whereas at 28°C they continued to increase. By 24 h.p.t. there was at least a 10-fold higher level of CHIKV RNA at 28°C compared to 37°C.

To confirm that this observation was not restricted to C2C12 cells, we also tested the replication of WT CHIKV-SGR-D-luc and the three mutants in Vero (African Green Monkey kidney epithelial) cells. Vero cells are defective in IFN α/β production due to a 9 Mb deletion in chromosome 12 but express the type 1 IFNAR1/2 receptor and can therefore respond to exogenous IFN [38, 39]. We again observed increased reporter activity of both WT and mutants in these cells at the lower temperature (Fig. S2E) suggesting that this temperature-driven difference in replication did not depend on intact IFN production.

We conclude that the three AUD mutants (W220A, D249A and Y324A) are temperature sensitive, exhibiting no defective phenotype in mammalian cells at a sub-physiological temperature (28°C). However, these experiments also revealed the novel observation that replication and/or translation of the WT CHIKV SGR was significantly enhanced at a sub-physiological temperature, and we sought to further investigate this intriguing phenomenon.

### Enhanced replication at sub-physiological temperature is not dependent on the reporter protein and is not observed for other arboviruses

As the CHIKV-SGR-D-Luc construct contains two reporters, we considered it prudent to confirm that the phenotype we observed was not influenced by these reporters, or by the insertion of a reporter into nsP3. To test this, two alternative CHIKV SGRs were utilized, namely CHIKV-SGR-nsP3-mCh-FF-Luc (in which the R-Luc reporter was replaced with mCherry) and CHIKV-SGR-FF-Luc (lacking R-Luc and thus expressing native nsP3) (Fig. S3A). These two SGRs were assayed in parallel with CHIKV-SGR-D-luc in C2C12 cells at the two temperatures (28°C or 37°C). All three showed enhanced levels of replication at 28°C compared to 37°C (Fig. S3B). As described previously (Fig. 1B), this effect was most apparent between 12-24 h.p.t. These data confirm that enhanced genome replication at sub-physiological temperature is not dependent on the nature of the nsP3-fused reporter protein.

We reasoned that the enhancement of genome replication at sub-physiological temperature might be a common feature in other arbovirus species that share with alphaviruses a mode of transmission from mosquitoes to humans via a peripheral bite. We therefore tested replication of mini-genome constructs derived from Bunyamwera virus (BUNV), the type species of the genus Orthobunyavirus (negative-sense, single-stranded RNA viruses) [40] and an SGR derived from Zika virus, a positive-sense, single-stranded RNA virus in the *Flaviviridae* family [41]. As shown in Fig. S3C/D, although both the BUNV mini-genome [42] and the Zika nano-luc SGR [43] replicated poorly in BSR-T7 and BHK-21 cells respectively, they exhibited similar levels of genome replication at 28°C or 37°C, indicating that this phenotype is not a common feature in other arboviruses.

To determine if the temperature sensitivity of the AUD mutants was unique to CHIKV or conserved within other alphaviruses, we generated the same panel of mutants in the context of an infectious clone of the closely related alphavirus, O’Nyong Nyong virus (ONNV). In this clone, the fluorescent reporter ZsGreen was expressed from a sgRNA transcribed from a second copy of the sub-genomic promoter (ONNV-2SG-ZsGreen) (Fig. S4A) (kind gift from Andres Merits, University of Tartu). This construct enabled rapid assessment of genome replication following transfection, by quantification of both ZsGreen fluorescence per cell and the number of ZsGreen positive cells. Although expression of ZsGreen was dependent on genome replication as production of the sgRNA is templated from the negative strand, the levels of ZsGreen expression are representative of both replication and translation of the sgRNA. As shown in Fig. S4B, the three AUD mutants (W220A, D249A and Y324A) showed a defect in replication at 37°C as judged by a reduction in the number of ZsGreen positive cells, whereas at 28°C the replication of all three was indistinguishable from WT. In addition, the absolute levels of ZsGreen per cell for WT was approximately 2-fold higher at 28°C compared to 37°C, consistent with enhanced genome replication at the sub-physiological temperature.

### AUD mutants remain temperature sensitive in the context of infectious CHIKV

As we had previously identified a novel function of the CHIKV AUD in promoting virus assembly, we wished to determine whether the phenotypes we had observed in the context of CHIKV SGR were also seen in the context of infectious virus. The three AUD mutations (W220A, D249A and Y324A) were therefore generated in the context of the ‘Integration of Chikungunya virus research’ (ICRES) CHIKV infectious clone, also derived from the ECSA strain LR2006 OPY1 isolate [34]. *In vitro* transcribed RNA for each construct was transfected into C2C12 cells, cells were grown at either 28°C or 37°C, and released virus was harvested at 48 h post infection (h.p.i.). All supernatants were titrated by plaque assay on BHK-21 cells grown at 32°C. As shown in Fig. 2A, the three mutants produced significantly less infectious virus than WT at 37°C (between 10^3^ and 10^4^ reduction in titre). At 28°C, unlike the SGR data where the mutants were indistinguishable from WT, the mutants exhibited significantly reduced titres although this phenotype was much less pronounced than at 37°C (between 5-60 fold reduction at 28°C). Sequence analysis of ICRES-CHIKV AUD mutants at 28°C or 37°C indicated that no reversion had occurred at either temperature (data not shown). Unlike the marked differences in genome replication observed using SGRs, there was only a modest increase (3-fold) in virus production at 28°C compared to 37°C. This suggested that the higher level of genome replication at 28°C did not lead to a concomitant increase in assembly and release of infectious virus, however, we noted that this result was obtained following transfection of *in vitro* transcribed RNA. We therefore performed a one-step growth curve analysis, evaluating virus production and genome replication over a time-course of 96 h following infection of C2C12 cells at a low MOI. As shown in Fig. 2B, the production of infectious virus was delayed at 28°C compared to 37°C, however, the maximal titre reached was 5-fold higher (1.8 x 10^8^ at 72 h.p.i. compared to 3.6 x 10^7^ at 24 h.p.i.). A similar delay was observed when levels of intracellular positive and negative strand CHIKV RNA were quantified (Fig. 2C). As there was no marked difference in the maximal level of positive strand RNA at both temperatures this may be due to a higher level of packaging of the positive strand RNA into nascent infectious virus and release from the cell at the lower temperature (Fig. 2B). In contrast, the level of negative strand RNA did attain a higher value. These data are consistent with the results obtained using SGRs, namely that at later time points after infection the rate of genome replication was limited at 37°C but this limitation was relieved at the lower temperature. However, it should be noted that the differences observed in the context of infectious virus were not as marked as those seen in the SGR experiments, perhaps reflecting other rate limiting cell factors or processes involved in assembly and release of infectious virus.

**Figure 2.**
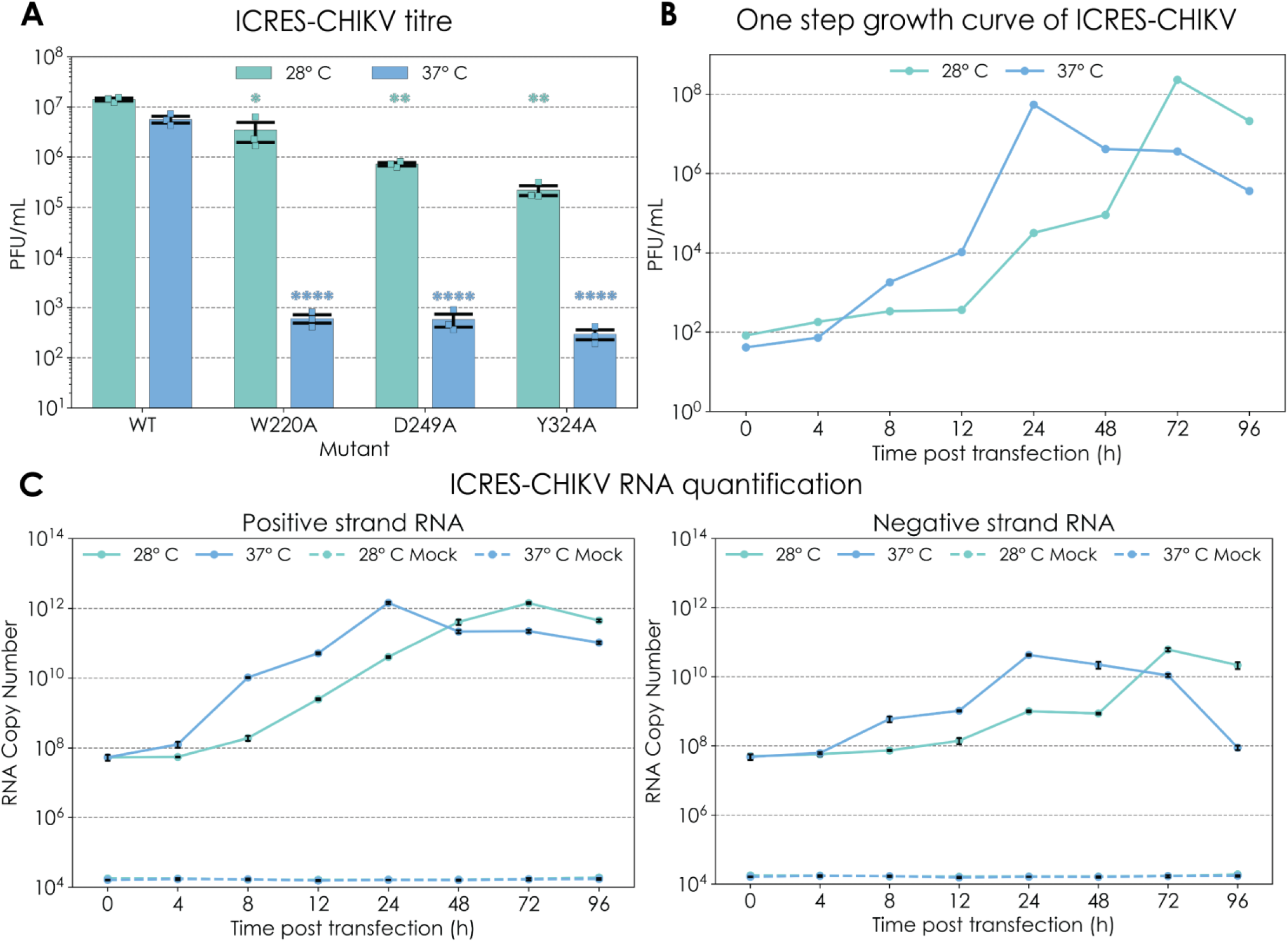
Phenotype of WT and AUD mutants in the context of ICRES-CHIKV at 28°C or 37°C. (**A**) Phenotype of WT and AUD mutants in ICRES-CHIKV production at 28°C or 37°C. C2C12 cells were transfected with WT and mutant ICRES-CHIKV RNA and incubated at 28°C or 37°C. Supernatants were collected 48 h post infection (h.p.i.) from both temperatures and titrated at 37°C by plaque assay. Significant differences denoted by * (P<0.05), ** (P<0.01) or **** (P<0.0001). Data are displayed as the means ±S.E. of three experimental replicates. (**B**) One step growth curve of ICRES-CHIKV in C2C12 cells at 28°C or 37°C. C2C12 cells were infected with ICRES-CHIKV at MOI 0.1 and incubated at 37°C or 28°C respectively. Cell supernatants were collected at indicated time points and virus titers were determined by plaque assay. (**C**) CHIKV RNA quantification from infected C2C12 cells. C2C12 cells collected at indicated time points were lysed by TRIzol solution, and total RNA were purified by phenol-chloroform extraction. cDNAs of CHIKV positive or negative strand RNA were synthesized as described before [27].

### Temperature shift assay reveals that early events in SGR-D-Luc replication are not enhanced at sub-physiological temperature

Previous studies have investigated temperature-sensitive mutants of alphaviruses by use of a temperature shift assay [44–46]. To further explore the role of temperature in controlling CHIKV genome replication, a temperature shift assay was conducted in C2C12 cells. WT CHIKV-SGR-D-Luc RNA was transfected into C2C12 cells and incubated either at 37°C or 28°C for 12 h, with and without a subsequent temperature shift from 12-24 h.p.t. (28°C to 37°C (designated S*), or 37°C to 28°C (designated S#)) (Fig. 3A). As seen previously (Fig. 1A), replication of WT CHIKV-SGR-D-Luc (as assessed by FF-Luc levels) up to 12 h.p.t. proceeded at broadly similar rates at both temperatures (Fig. 3B), and between 12-24 h.p.t. replication increased ∼100-fold at 28°C but did not increase at 37°C. However, when the temperature was shifted from 28°C to 37°C at 12 h.p.t. (S*), the replication increased only ∼20-fold between 12-24 h.p.t. (Fig. 3B, left). The opposite effect was seen when transfected cells were first incubated at 37°C: replication reduced slightly between 12-24 h.p.t. at 37°C, however, when the temperature was shifted from 37°C to 28°C at 12 h.p.t., replication increased 5-fold between 12-24 h.p.t. (Fig. 3B, right).

**Figure 3.**
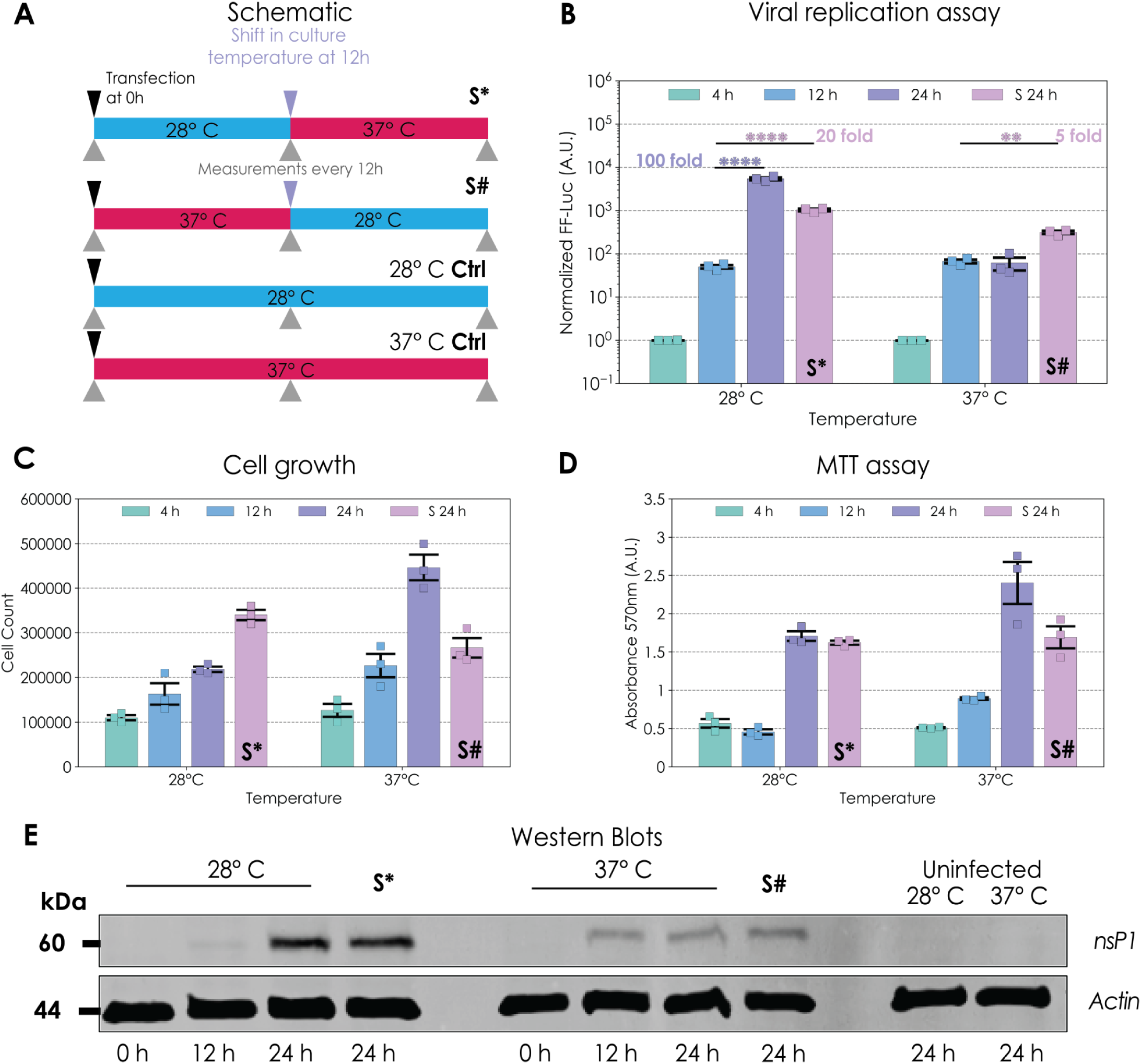
Temperature shift assay of WT SGR-D-Luc. (**A**) Schematic of temperature shift assay. Transfected C2C12 cells were incubated at either 28°C or 37°C prior to a temperature shift from 12-24 h.p.t. (28°C to 37°C (designated 24 h S*), or 37°C to 28°C (designated 24 h S#). Controls maintained the same temperature from 0-24 h.p.t. (**B**) Replication assay. WT CHIKV-SGR-D-Luc RNA was transfected into C2C12 cells and incubated at 28°C or 37°C according to the schematic in **(A)**. FF-Luc activities were measured and normalized to their respective 4 h values. Significant differences denoted by ** (P<0.01) or **** (P<0.0001), compared to replication at 12 h.p.t. Data are displayed as the means ±S.E. of three experimental replicates. Cell numbers were enumerated (**C**) and cell metabolic activity **(D)** determined by MTT assay at 28°C or 37°C. **(E)** Protein expression of nsP1 at 28°C or 37°C. Cells were lysed at 24 h.p.t. and 10 µg of each sample was analysed by western blot. Actin was also detected by western blot as a loading control.

Importantly, both cell growth assessed by quantification of viable cells (Fig. 3C), and metabolic activity assessed by MTT assay (Fig. 3D) were higher at 37°C compared to 28°C. Temperature shifts elicited the expected changes in both cell number and metabolic activity. These data confirmed that the enhanced replication observed at sub-physiological temperature cannot be explained by increased cell growth and metabolic activity. In fact, we observed an inverse correlation between virus genome replication and cell growth and metabolic activity. These data were also confirmed by western blotting for nsP1 (Fig. 3E), demonstrating that translation of the 5’ ORF was also enhanced at the sub-physiological temperature.

### Enhanced genome replication at a sub-physiological temperature is correlated with impaired expression of ISGs

We considered that one reason for the increased level of CHIKV genome replication at 28°C might be due to a sub-optimal antiviral transcriptional response, even in the absence of exogenous IFN stimulation. It has previously been documented that IFN responses are impaired at sub-physiological temperatures, and this has been proposed to explain the enhanced CHIKV replication and pathogenicity observed in mice housed at 22°C compared to 30°C [31].

We first asked if the transcriptional response of C2C12 cells to stimulation with the synthetic dsRNA analogue poly I:C was affected by temperature. C2C12 cells at either 28°C or 37°C were mock treated or stimulated with poly I:C for 24 h prior to analysis of the transcription of three well characterized ISGs by qRT-PCR. As shown in Fig 4A, at 37°C poly I:C treatment significantly enhanced the transcription of *Ifi44i*, *Cmpk2* and *Uba7* compared to cells treated at 28°C, or mock treated cells. This is consistent with impairment of the ISG response at 28°C.

**Figure 4.**
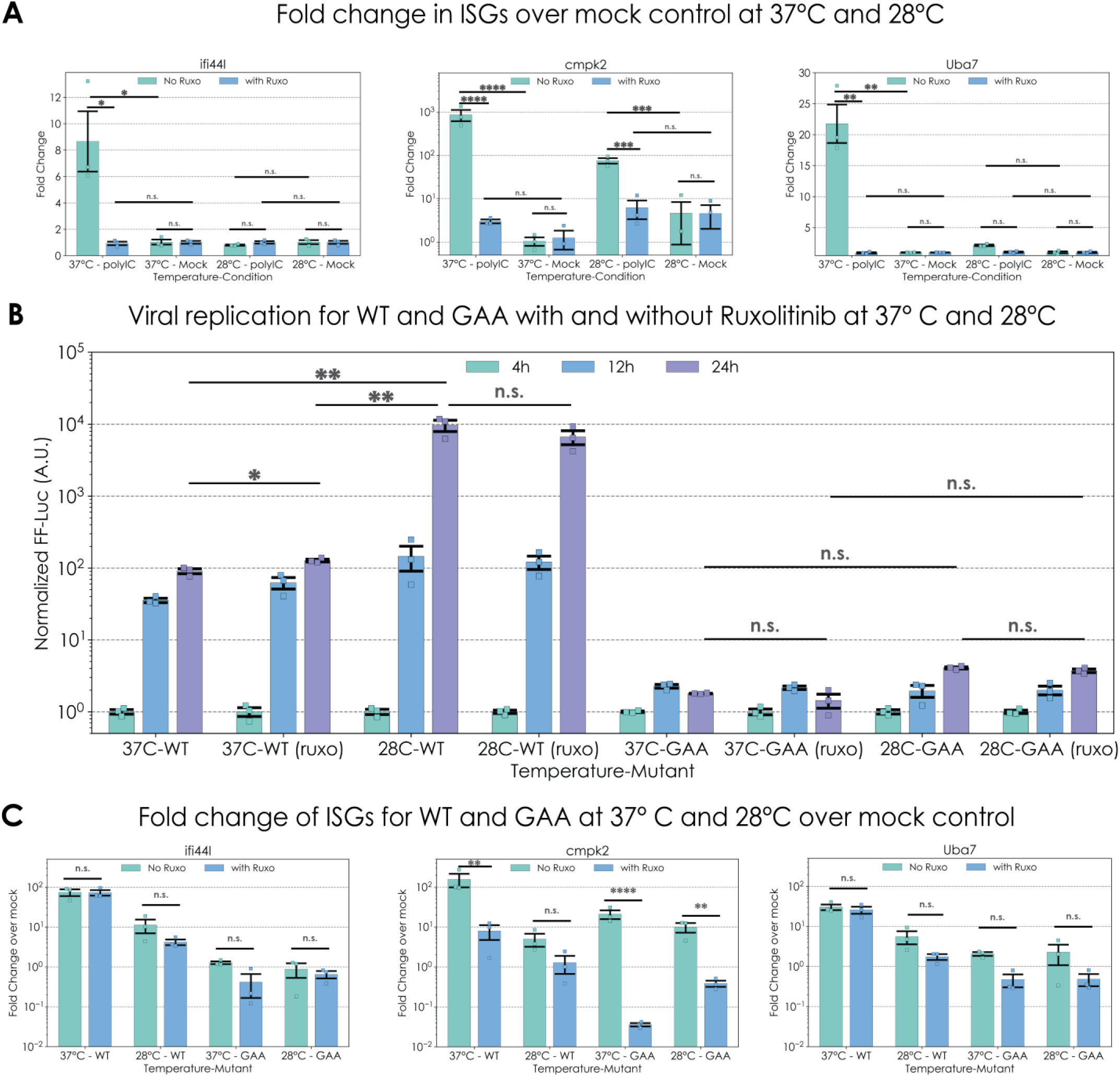
Effect of ruxolitinib (Ruxo) treatment on ISG expression in cells transfected with poly I:C or WT CHIKV-SGR-D-Luc. (**A**) Effect of Ruxo on ISG expression in response to poly I:C treatment. C2C12 cells in 24 well plates were pre-treated with 10 uM Ruxo for 12 h before transfection with 1 ug poly I:C per well and ISG expression was measured by qRT-PCR at 24 hpt. (**B**) Replication of WT and GAA CHIKV-SGR-D-Luc in response to Ruxo at 28°C or 37°C. C2C12 cells were first pre-treated with 10 uM Ruxo 12 h before transfection of WT and GAA CHIKV-SGR-D-Luc. Cells were then incubated at 37°C or 28°C and harvested at 4, 12 and 24 hpt. (**C**) Fold change analysis of ISGs for WT and GAA at 37°C or 28°C compared to mock control. Fifty µl of lysed samples was subjected to RNA extraction, followed by elution in RNase-free H₂O. Ct values for three ISGs were measured alongside the housekeeping gene β-actin across different samples, and fold changes were calculated accordingly. Significant differences denoted by n.s. (P>=0.05), * (P<0.05) or ** (P<0.001) compared to WT are only shown for the 24 h time point for clarity. Data are displayed as the means ±S.E. of three experimental replicates.

To test whether this poly I:C response involved canonical IFN signalling via the JAK-STAT pathway, we treated cells with ruxolitinib (Ruxo: a JAK-STAT inhibitor). We first determined that the maximal non-cytotoxic concentration of Ruxo in C2C12 cells was 10 µM (Fig S5). As expected, treatment of cells with 10 µM Ruxo blocked the poly I:C stimulated transcription of these 3 genes (Fig 4A), suggesting that this was acting via the JAK-STAT pathway presumably in an autocrine fashion following the poly I:C stimulation of IFN expression.

We next tested whether there was a temperature sensitive effect of JAK-STAT signalling on CHIKV replication. Specifically, whether inhibition of JAK-STAT signalling at 37°C could relieve the block to genome replication between 12-24 h. However, as shown in Fig 4B, Ruxo treatment had no significant effect on the replication of the CHIKV WT D-luc SGR at either temperature. We also examined the effect of active CHIKV genome replication on the transcription of the 3 known ISGs: *Ifi44l*[47], *Cmpk2* [48], and *Uba7* [49]. As shown in Fig 4C, induction of ISG transcription by CHIKV was higher at 37°C than at 28°C. Notably, with the exception of *Cmpk2*, the transcriptional upregulation was not affected by Ruxo treatment. Taken together these data indicate that CHIKV replication induced much higher levels of these 3 ISGs at 37°C than at 28°C, and that unusually this ISG induction was not halted by pharmacologic inhibition of JAK-STAT signalling, hinting at non-canonical ISG activation.

Therefore, we next wished to determine whether only these 3 ISGs were induced, or whether a global ISG induction also occurred during CHIKV infection. To test this, we conducted a global transcriptomic analysis. We performed bulk RNA-Seq on total RNA from C2C12 cells transfected with either WT CHIKV-SGR-D-luc, or the non-replicative GAA mutant, harvested at either 12 h or 24 h.p.t. We performed pairwise differential gene expression analysis at the two temperatures, or between WT and GAA transfected cells. As expected, this analysis revealed many differentially expressed genes (defined as p adj. < 0.05, and [log2 fold change] >1) both in response to CHIKV genome replication, but also between the two temperatures.

In this study, to assess effects on CHIKV genome replication, we focused our analysis on known ISGs (annotated from the Molecular Signatures database (MSigDB) HALLMARK_INTERFERON_ALPHA_RESPONSE dataset) [50]. In agreement with our prior RT-qPCR data (Fig. 4C), at 12 h.p.t., *Ifi44l*, *Cmpk2*, and *Uba7* were upregulated in WT- versus GAA-transfected cells cultured at 37°C but not at 28°C, along with a number of other prominent ISGs, such as *Batf2, Cxcl11,* and *Oasl1*, as well as immunoproteosome components such as *Psmb8* (Figs. 5A-B). Across all 94 annotated ISGs, this amounted to 32 ISGs that were potently (up to 512-fold) up-regulated at 37°C, versus only 2 ISGs (*Ifit3*, *Ccrl2*) that were modestly (up to 8-fold) up-regulated at 28°C (Figs. 5A-D, 6). More ISGs were up-regulated at 24 h.p.t. in comparison to 12 h.p.t at 37°C or 28°C; nevertheless we observed the same trend: both the number of ISGs, and the level of upregulation, were markedly greater at 37°C compared to 28°C at 24 h.p.t. (Figs. 5E-H and 7). At 37°C, 52 out of 94 ISGs were up-regulated but only 11 of these ISGs were up-regulated at 28°C (Fig. 7). When we ranked the fold-changes of gene expressions between WT-transfected cells at 37°C versus 28°C, the 94 annotated ISGs were strongly enriched among the highest ranks (Fig. 5I, Normalized enrichment score (NES) -2.09, q-value 0.00 at 12 h.p.t.; Fig. 5J, NES -2.85, q-value 0.00 at 12 h.p.t.), but not when the replication-deficient GAA was transfected (in this case the data failed to generate a valid score, likely due to insufficient enrichment or technical limitations in the dataset, data not shown). To validate these data we undertook qRT-PCR for 6 selected genes (*Uba7, Stat2, Ifi44I, Isg20, Oas1a* and *Cmpk2*) for which well-characterised primers were available in house. As shown in Fig. S6 the overall levels of these 6 genes were higher at 37°C compared to 28°C, at either 12 h.p.t or 24 h.p.t. Overall, these data support the conclusion that global ISG expression in response to CHIKV genomic replication is significantly impaired at the sub-physiological temperature.

**Figure 5.**
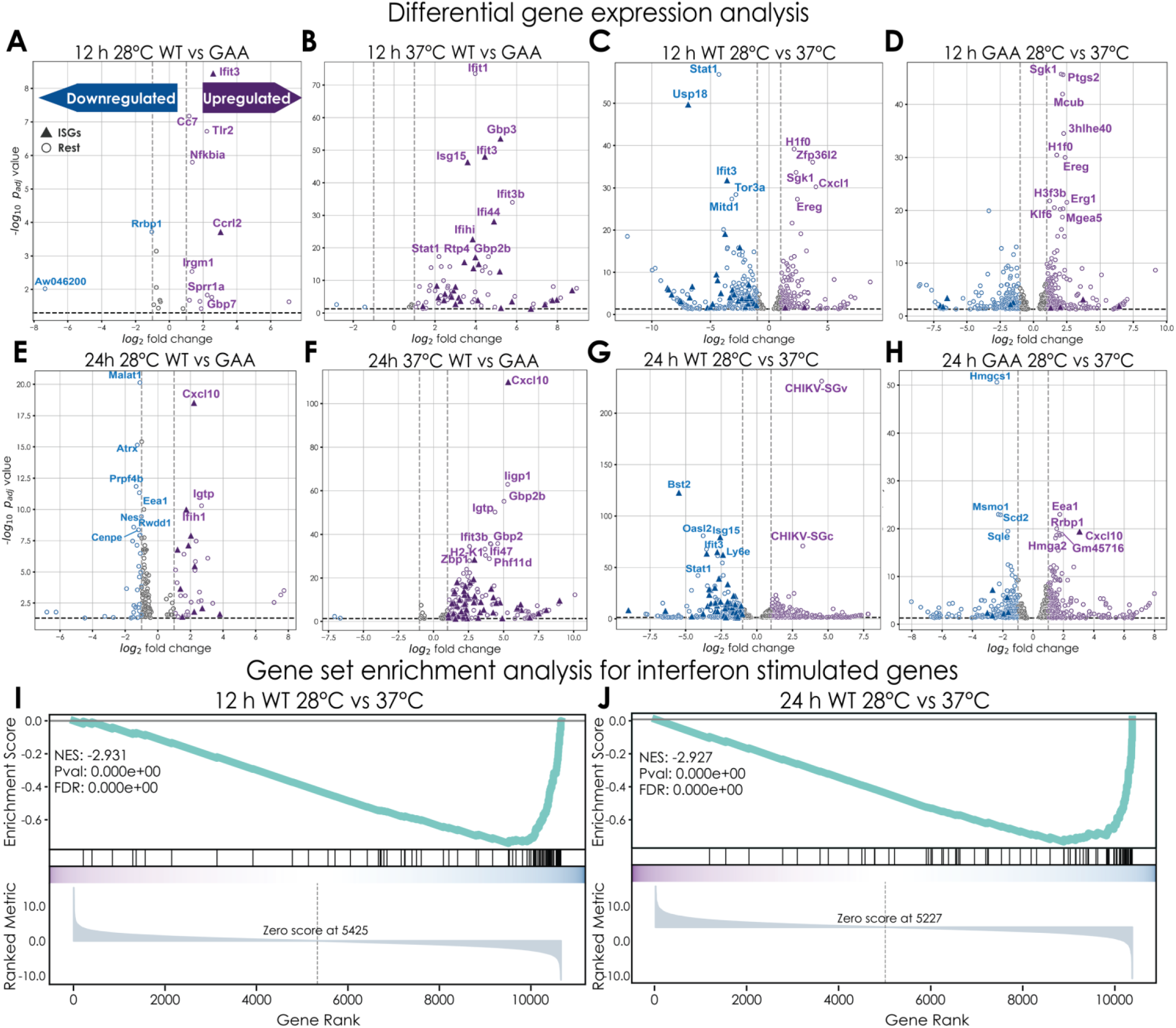
Volcano plots of differential expressed genes in response to WT CHIKV-SGR-D-Luc replication at 28°C or 37°C. C2C12 cells were transfected with WT CHIKV-SGR-D-Luc at 28°C or 37°C as described above. At 12 h.p.t or 24 h.p.t., total RNA was harvested and subjected to bulk RNA-Seq. (**A-H**) Volcano plots of pairwise differential gene expression between the different conditions. The IFN-stimulated genes (ISGs) that were significantly expressed are labeled as triangles while non-ISGs are labeled as dots. (**I and J**) Waterfall plots of gene set enrichment analysis (GSEA) of differentially expressed genes within ISGs. ISGs were defined as the 94 genes within the *M. musculus* Molecular Signatures Database (MSigDB) HALLMARK_ALPHA_RESPONSE [50].

**Figure 6.**
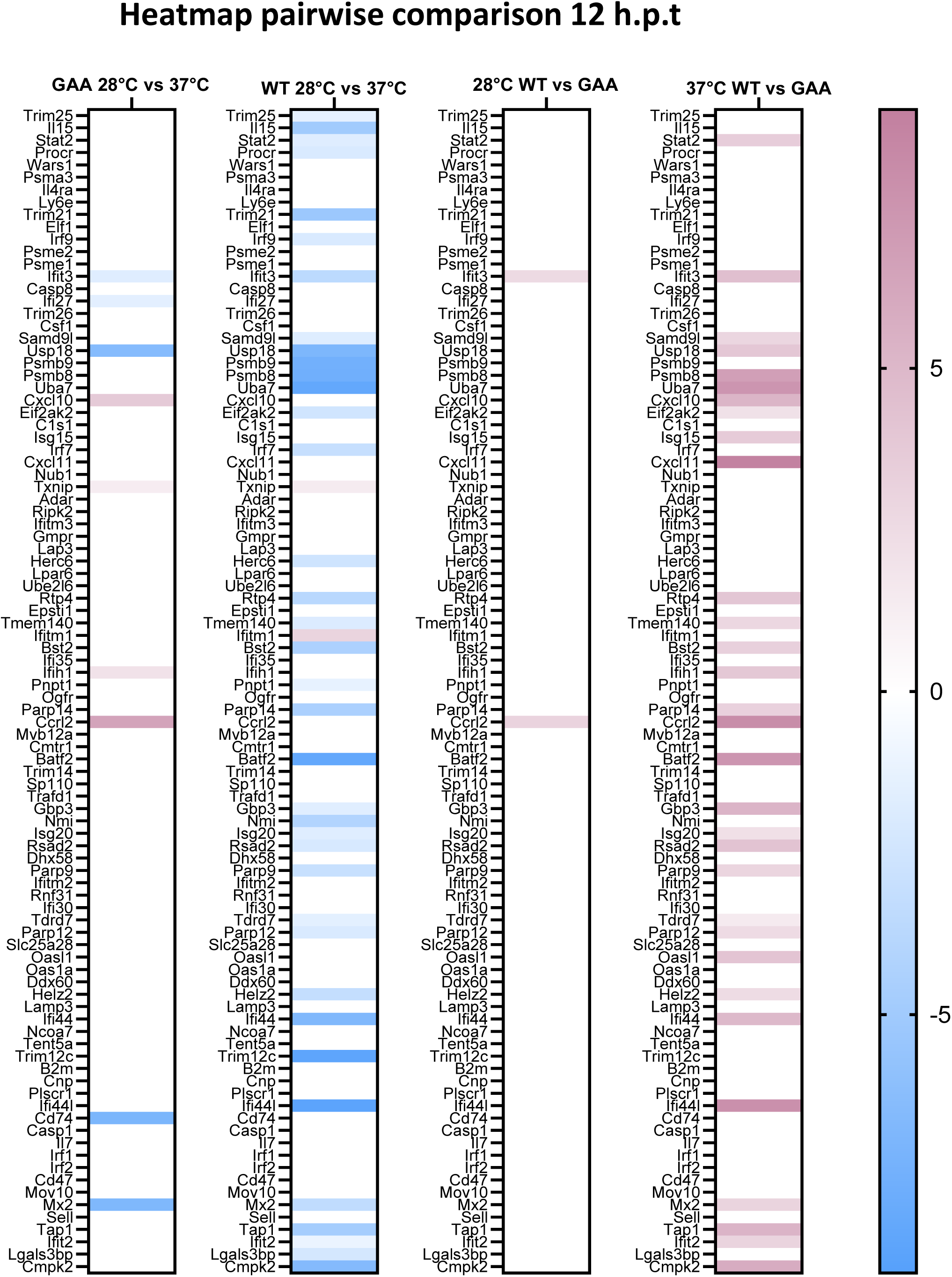
Heatmap of ISGs in response to WT CHIKV-SGR-D-Luc replication at 28°C or 37°C at 12 h.p.t. The differentially expressed ISGs from the volcano plots at 12 h.p.t. were selected and grouped by R-studio. The heatmap plot was generated using Graphpad Prism version 9.0.

**Figure 7.**
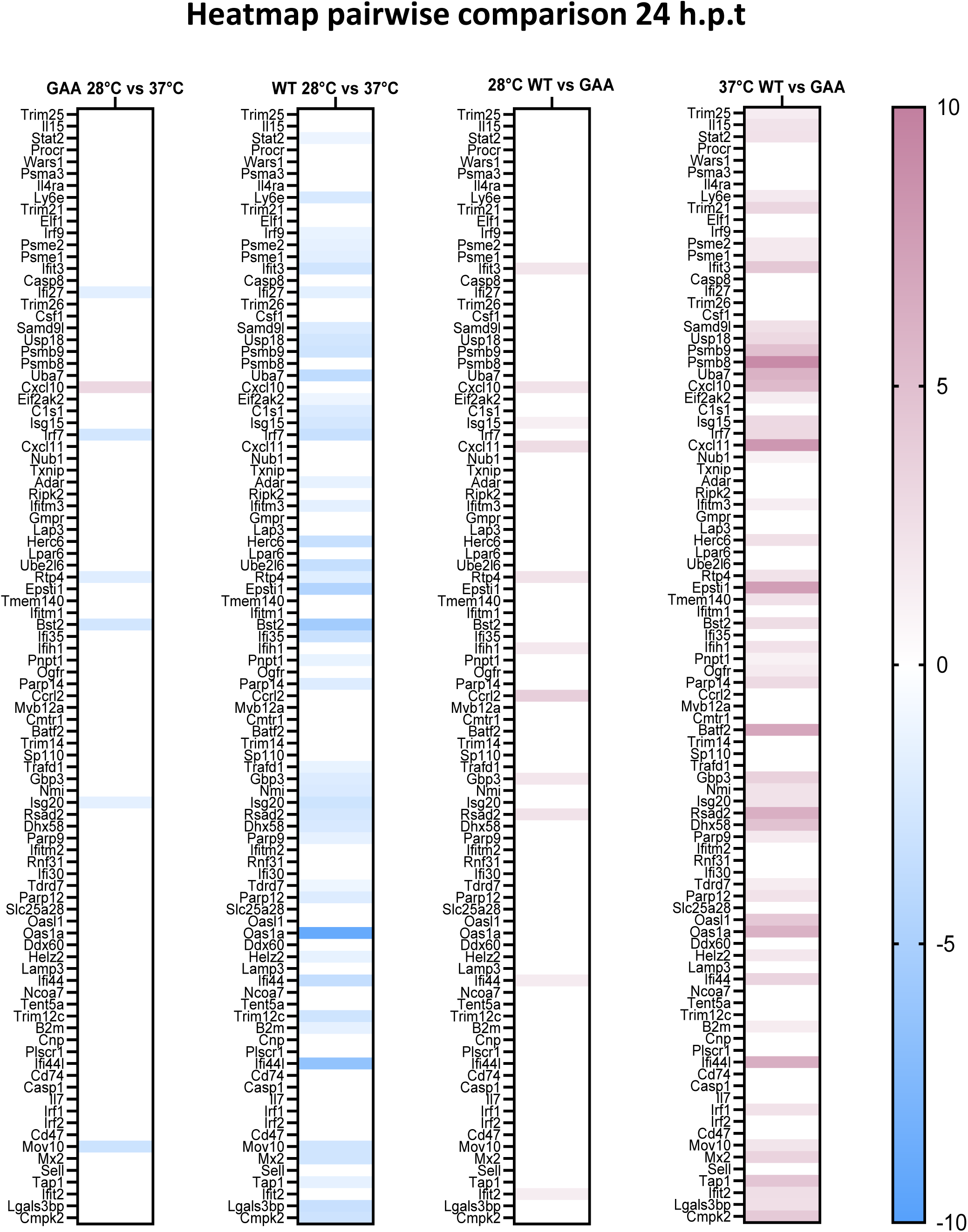
Heatmap of ISGs in response to WT CHIKV-SGR-D-Luc replication at 28°C or 37°C at 24 h.p.t. The differentially expressed ISGs from the volcano plots at 24 h.p.t. were selected and grouped by R-studio. The heatmap plot was generated using Graphpad Prism version 9.0.

### Enhanced recruitment of the key SG protein G3BP1 to replication sites at a sub-physiological temperature

In addition to the ISG response, other cellular pathways may act in an antiviral fashion. Cell stress triggers SG assembly, which halts translation and therefore may be detrimental to viral replication. During alphavirus infection, nsP3 sequesters key SG proteins such as G3BP1/2, and recruits them to cytoplasmic foci where viral genome replication occurs, leading to inhibition of the formation of *bona fide* SG [23–25, 51]. We considered that this function of nsP3 might play a role in the enhanced replication observed in mammalian cells at a sub-physiological temperature. SG formation can be induced by treatment with sodium arsenite (NaAsO_2_) which results in oxidative stress, and they can be disassembled by cycloheximide (CHX) treatment.

Importantly, it has been previously shown that SG formation may be temperature-dependent, as NaAsO_2_ was only able to induce SG at 37°C but not 28°C [52]. Using G3BP1 (detected using a pan-G3BP antibody) as a marker for the formation of bona fide SG we confirmed that this was also the case in our hands (Fig. S7), as no G3BP1-positive SG were observed following NaAsO_2_ treatment of C2C12 cells at 28°C. As expected, at 37°C SG were efficiently induced by NaAsO_2_ treatment and disassembled by CHX (Fig. S7).

To investigate this further, we exploited the CHIKV SGR-nsP3-mCh-FF-luc construct in which nsP3 was fused to mCherry (see Fig. S3A), and we analysed the distribution and abundance of nsP3-mCherry and G3BP1 in transfected C2C12 cells (Fig. 8). As a further marker for sites of active genome replication (cytoplasmic foci) we also stained cells with an antibody to visualize dsRNA (J2), previously shown by us [1] and others [24, 53] to colocalise with nsP1, nsP3 and G3BP1. Consistent with previous data in this manuscript (eg Fig. S3B), at 24 h.p.t., both dsRNA and nsP3-mCherry accumulated to a higher level at 28°C compared to 37°C, and this enhancement was not seen at earlier timepoints (Fig.8B). In contrast, the amount of G3BP1 in transfected cells increased over time and was higher at 37°C when compared to 28°C (Fig. 8B).

**Figure 8.**
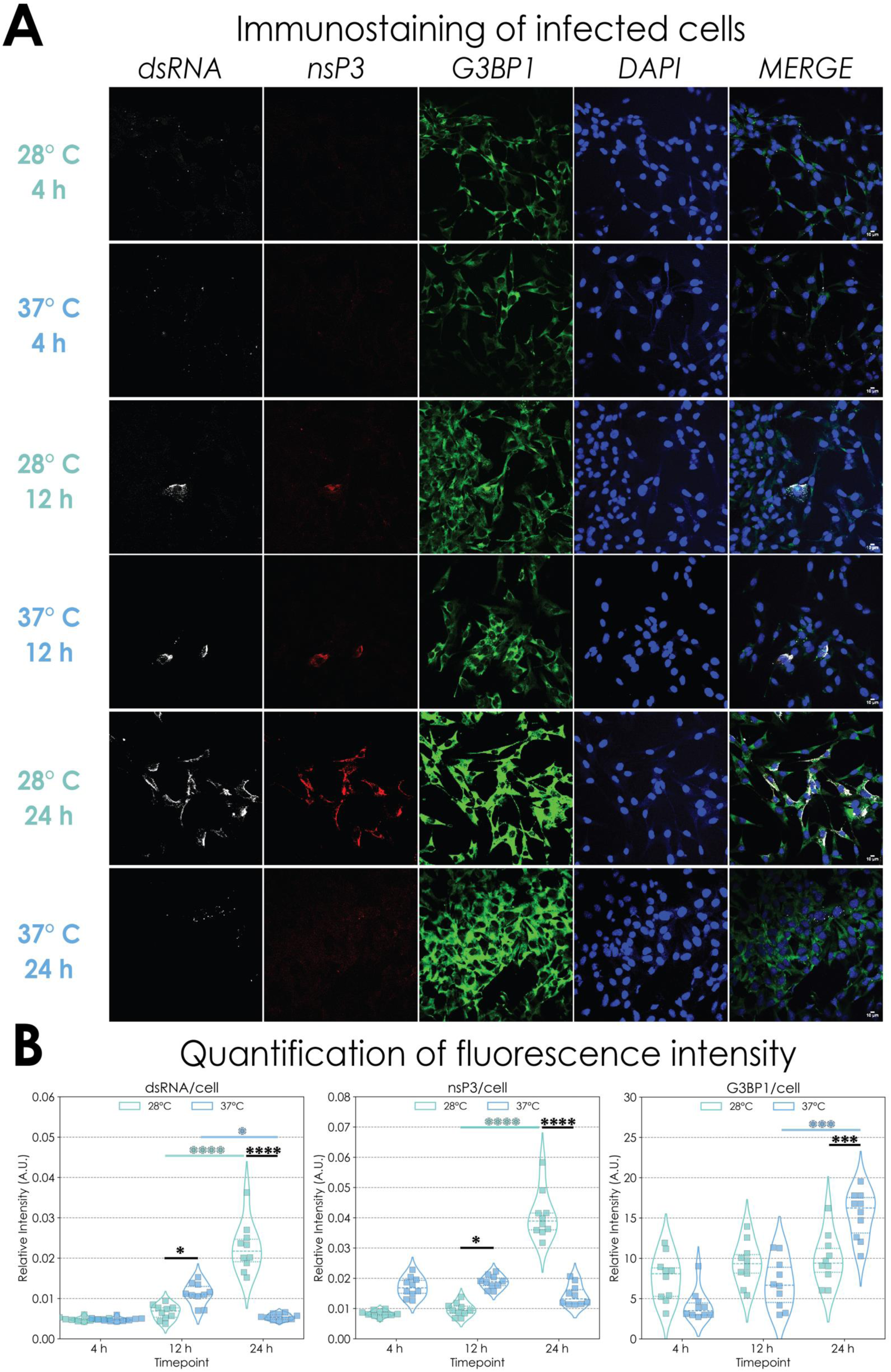
Formation of cytoplasmic nsP3/dsRNA positive foci at 28°C or 37°C. (**A**) C2C12 cells were transfected with WT CHIKV-SGR-nsP3-mCh-FF-Luc and incubated at 28°C or 37°C respectively. Cells were then fixed with 4% paraformaldehyde and stained with antibodies to G3BP1 (Green) or dsRNA (gray). nsP3-mCherry was expressed as a fused protein (Red). Scale bar 10 µm. (**B**) Quantification of the overall dsRNA, nsP3-mCherry and G3BP1 expression at 28°C or 37°C. Protein expression levels were determined for 10 cells and calculated by Fiji ImageJ. Significant differences denoted by * (P<0.05), *** (P<0.001) and **** (P<0.0001), compared to values at 37°C.

We then quantified the colocalisation of nsP3, G3BP1 and dsRNA (Fig. 9A). Quantitative analysis (Fig. 9B) showed that at 12 h.p.t. there were no differences in the amount of colocalized nsP3/G3BP1, nsP3/dsRNA or G3BP1/dsRNA between the two temperatures, however, at 24 h.p.t., the levels of colocalisation for all three pairs were significantly higher at 28°C compared to 37°C. Taken together, these data demonstrate that there are more replication sites, higher levels of dsRNA, and more recruitment of the SG protein G3BP1 to replication sites at 28°C than at 37°C.

**Figure 9.**
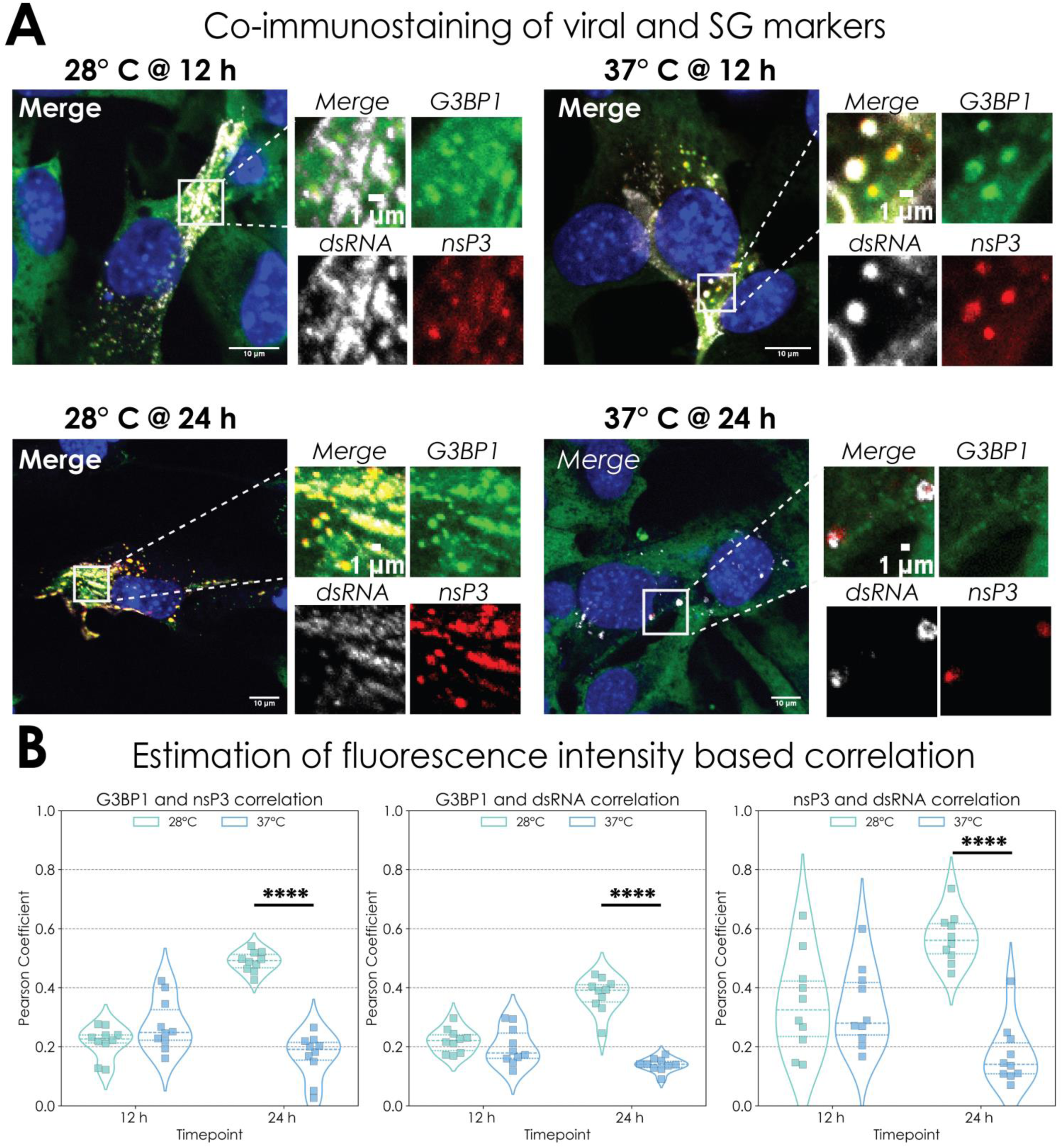
Colocalisation of G3BP1 with nsP3 and dsRNA at 28°C or 37°C. (**A**) Detection of dsRNA, nsP3 and G3BP1 in WT CHIKV-SGR-nsP3-mCh-FF-Luc transfected C2C12 cells. Immunofluorescence assays were performed as before. Colocalisation of dsRNA, nsP3 and G3BP1 is shown. All image analysis and processing was performed using Fiji software. Fluorescence plot profile of nsP3, dsRNA and G3BP1 signal intensities along the indicated white line is shown. (**B**) Quantitative analysis of the colocalization between dsRNA, nsP3 and G3BP1 signals. Pearson’s correlation coefficient between dsRNA and G3BP1, nsP3 and G3BP1, dsRNA and nsP3 were determined for 10 cells and calculated by Fiji ImageJ. Significant differences denoted by **** (P<0.0001), compared to values at 37°C.

## Discussion

This study set out to build on prior work regarding the functions of the CHIKV AUD [1], and analyzed the phenotype of a panel of alanine substitutions. Our study took an intriguing turn when we observed that 3 of these mutants (W220A, D249A and Y324A) exhibited a defective replication phenotype in mammalian cells at 37°C, but no defect in either mosquito cells, or in mammalian cells at a sub-physiological temperature (28°C), defining these as temperature-sensitive (ts) mutants (Figs 1B and S2B/C). Although ts mutations in alphavirus nsP3 have been previously identified, for example F312S and A268V in Sindbis virus, these resulted in a defect in negative-strand RNA synthesis at the non-permissive temperature (40°C) [54]. These data point to a critical role of nsP3, and the AUD in particular, in genome replication. Further biochemical analysis of these ts mutants in nsP3 may shed light on the specific role of nsP3 but is not the focus of this manuscript.

This analysis led us to the serendipitous discovery that wildtype CHIKV SGR replicated more efficiently at sub-physiological temperatures in mammalian cells. The novelty of this intriguing and arguably counter-intuitive observation led us to focus our attention on the underlying mechanisms. We note that, although a 100-fold enhancement of CHIKV genome replication at a sub-physiological temperature was observed in the context of CHIKV SGRs (Fig. 1B), the enhancement in the context of infectious virus was less dramatic (Fig. 2A). We observed only a modest (5-fold), but reproducible increase in virus production with delayed kinetics at sub-physiological temperatures, suggesting that other processes during the virus lifecycle (e.g. virus assembly and release) are also affected by temperature. This is consistent with the modest increase in maximum virus titre in mice infected with CHIKV sub-cutaneously through the feet and housed at 22°C compared to 30°C [31].

In considering why CHIKV genome replication in mammalian cells might be enhanced at sub-physiological temperatures, we reasoned that the first cells infected in a mammalian host via a mosquito bite would be in the periphery and thus not at 37°C. It would be reasonable to expect that other arboviruses would also exhibit the same differential, however, that was not the case as both BUNV and ZIKV replicated to similar levels at 28°C and 37°C (Fig S3C/D) This suggests that this phenotype is driven by alphavirus specific processes, and we note that neither the nsP3 macro-domain nor the AUD have functional homologues in either bunyaviruses or flaviviruses. Analysis of the timescale of genome replication at different temperatures also revealed that the phenotype was most apparent after 12 h.p.t. In most cases, replication rates up to 12 h.p.t. did not differ between the two temperatures (e.g. see Fig. 1A/B), however between 12-24 h.p.t. replication continued to increase at 28°C, but plateaued at 37°C.

An explanation for our observation was that it reflected an aspect of the cellular response to virus genome replication that was perhaps impaired at the sub-physiological temperature and only manifest after 12 h.p.t. An obvious candidate would be the type I IFN response, which has indeed been shown to be reduced at lower temperatures promoting CHIKV arthritis in a mouse model [31]. Consistent with this, the response to the synthetic dsRNA analog poly I:C was impaired at 28°C and blocked by the JAK-STAT inhibitor Ruxo. Similarly, the ISG transcriptional response to CHIKV genome replication (in SGR transfected cells) was severely impaired at the sub-physiological temperature (Figs. 5-7). At the sub-physiological temperature only a few ISGs from the total of 94 were up-regulated, while at 37°C many more ISGs were up-regulated (Fig 5C, D, G and H, and Fig 6). This data is also consistent with previous research [30], in which transcription of ISGs following IFN treatment was reduced at a sub-physiological temperature.

The exact antiviral mechanisms of ISGs against CHIKV, and whether reduced ISG expression fully explains temperature-dependent replication remains unclear. Here we attempted to study the antiviral effect of ISGs in two systems. First, we employed Vero cells, which cannot produce IFNα/β, and therefore we would expect these cells to have impaired ISG responses. In Vero cells, we found that CHIKV still replicates better at 28°C than at 37°C, suggesting that intact type I IFN production is not required for temperature-driven differences in replication. Second, we treated mouse C2C12 cells with Ruxo to inhibit JAK-STAT signaling (which is triggered by IFN signaling) and to thereby block ISG induction. Under canonical conditions, IFN production triggers a well-characterized JAK1/2-dependent phosphorylation cascade, which can be inhibited by Ruxo. This phosphorylation leads to the formation of ISGF3 (p-STAT1, p-STAT2, and IRF9), which subsequently binds to ISG promoters to activate transcription at later stages. However, under non-canonical conditions [55], antiviral responses can still be mediated by non-phosphorylated forms of these factors or alternative transcriptional regulators, bypassing JAK-STAT activation.

One key player in non-canonical signaling is IRF3, a major regulator of type I IFN production, which can also induce a subset of ISGs independently of IFN signaling. For example, IRF3 activation upregulates ISG54, ISG20, and IFP35 [56]. Similarly, during influenza virus infection, several ISGs, are induced without IFN production, but relies on activation of IRF3 and transcription induction of p56 gene [57]. Likewise, HCV infection triggers direct recruitment of NF-κB and IRF3 to the CXCL10 promoter via RIG-I and TLR3 sensing, bypassing type I/III IFN pathways [58]. In HEV infection, RIG-I plays a dual role, activating IRF3 and IRF7 via JAK-STAT pathway, while also inducing ISGs such as IFIT1, RANTES, and CXCL10 independently of STAT1 phosphorylation [59]. Similarly, during Coxsackievirus B3 infection, MDA5, rather than RIG-I, drives an IFN-independent antiviral response in inhibiting viral replication [60].

More recently, studies on CHIKV infection suggest that CHIKV actively subverts canonical IFN responses, enhancing its reliance on non-canonical signaling pathways to restrict viral replication [61]. These observations highlight the critical role of both canonical and non-canonical IFN signaling in antiviral immunity. A deeper understanding of these mechanisms could provide novel therapeutic strategies against CHIKV and other viral infections.

An alternate hypothesis is that reduced ISG expression is only one factor that allows for increased CHIKV replication at reduced temperatures, and other mechanisms may play roles in the phenotype. To further investigate the mechanisms underpinning the enhanced CHIKV replication at the sub-physiological temperature we analysed the production of SGs, cytoplasmic protein-mRNA aggregates formed in response to stress and believed to play an antiviral role [18]. Alphavirus infection has previously been documented to disrupt formation of *bona fide* SG by recruiting key SG components such as G3BP1/2 to cytoplasmic foci where genome replication occurs [24], as judged by positivity for both nsP3 and dsRNA. G3BP1/2 have been shown to be critical for alphavirus genome replication [21]. Sequestration of G3BP1/2 away from *bona fide* SG will have the added benefit of limiting the antiviral effects of SG [18].

Interestingly, as seen previously [52], treatment with NaAsO_2_ (which induces oxidative stress) did not result in SG formation at 28°C (Fig S6), suggesting that there was a defect in this process at the sub-physiological temperature. However, following transfection with a CHIKV SGR (expressing an mCherry-nsP3 fusion) we observed that, whereas total levels of G3BP1 were higher at 37°C (Fig 8), it was more efficiently recruited into nsP3/dsRNA positive cytoplasmic foci at 28°C (Fig 9). Specifically, at 24 h.p.t. not only were there more nsP3/dsRNA positive cytoplasmic foci at 28°C (Fig. 8), but also the recruitment of G3BP to nsP3/dsRNA foci was significantly higher at 28°C (Fig 9).

These data are consistent with the hypothesis that the enhanced level of CHIKV genome replication in mammalian cells at a sub-physiological temperature is associated with more efficient recruitment of cellular factors to cytoplasmic sites of replication. A likely explanation is that the absence of *bona fide* SG formation at 28°C (Fig. S7) means that SG components are more abundant in the cytoplasm, and this facilitates recruitment by nsP3 (or other nsPs). In addition, our data suggest that the availability of SG components is not a limiting factor for genome replication until after 12 h.p.t. as there are no differences in replication or recruitment of G3BP1 to cytoplasmic foci at this timepoint at either temperature (see Figs. 1A, 9B).

Why the process of recruiting SG components to cytoplasmic sites of replication should differ at different temperatures remains to be determined. Importantly the ADP-ribosylhydrolase activity of the nsP3 macro-domain has recently been shown to play a key role in SG formation, disassembly and composition [25]. It is possible that the enzymatic activity and/or substrate specificity of the macro-domain could be modulated by temperature. Although the AUD *per se* has not been directly shown to play a role in SG formation, it is flanked by the macro-domain and HVD, both of which have now been implicated in SG formation. Localization of CHIKV nsP3 to cytoplasmic foci in invertebrate cells was shown to depend on both the AUD and the HVD [62], although it should also be noted that the interaction with G3BP1/2 (or its insect homologue, Rasputin) involves specific sequences within the HVD [63]. It is thus possible that all three domains of nsP3 coordinate to control formation of *bona fide* SG, and recruit SG components to cytoplasmic sites of genome replication.

In conclusion, our data have revealed a hitherto unrecognized enhancement of CHIKV genome replication in mammalian cells at sub-physiological temperature that may play a role in the early stages of transmission of virus to the mammalian host via a mosquito bite. Our observations point to a critical role of the AUD in this difference, and further work seeking to understand the effects of temperature on nsp3 interactions with host cell proteins would benefit greatly from a proteomic approach.

## Materials and Methods

### Sequence alignment

AUD amino acid sequences from a selection of alphaviruses (Sindbis, Semliki, Ross River, Venezuelan, Eastern and Western Equine Encephalitis, ONNV and CHIKV) were obtained from NCBI and aligned by Clustal Omega. The conserved AUD residues were identified by Pymol and their locations were predicted from the Sindbis nsP2/3 protein structure (PDB ID code 4GUA) [32].

### Cell culture

Vero and BHK-21 cells were grown in DMEM supplied with 10% foetal calf serum (FCS), 0.5 mM non-essential amino acids and penicillin-streptomycin (100 units/mL). C2C12 cells were grown as above but with 20% FCS. All mammalian cells were maintained at 37°C or 28°C with 5% CO_2_. BHK-21 cells were obtained from John Barr and C2C12 cells from Michelle Peckham (University of Leeds). U4.4 and C6/36 cell lines (*Ae. albopictus*) were originally obtained from Susan Jacobs (The Pirbright Institute) and were cultured in Leibovitz’s L-15 media containing 10% FCS, 10% tryptose phosphate broth and penicillin-streptomycin (100 units/mL), at 28°C without CO_2_. Infection and viral RNA transfection experiments were carried in accordance with protocols reviewed and approved by the Institutional Biosafety Committees of the University of Leeds and Princeton University (# 1175)

### Plasmid constructs

CHIKV-SGR-D-Luc was used as a template for site-directed mutagenesis using specific primers to generate each of the AUD mutations. The AUD mutated fragments were excised and ligated back into the CHIKV-SGR-D-Luc backbone, or into the infectious ICRES-CHIKV clone [34]. For construction of the ONNV AUD mutants, a fragment including the AUD coding sequence was excised from the ONNV-2SG-ZsGreen (donated by Andres Merits, University of Tartu), and ligated into pcDNA3.1 to generate pcDNA3.1-ONNV-AUD. This was used as a template for site-directed mutagenesis using specific primers to generate ONNV AUD mutations. The ONNV AUD mutated fragments were excised from pcDNA3.1-ONNV-AUD and ligated into the ONNV-2SG-ZsGreen backbone. CHIKV-SGR-nsP3-mCh-FF-Luc was generated by SpeI digestion of the FF-Luc fragment from SGR-D-Luc and insertion of the mCherry coding sequence. CHIKV-SGR-FF-Luc was constructed by SpeI digestion of the R-Luc fragment from SGR-D-Luc and re-ligation.

### Transfection and dual luciferase assay

Capped RNAs of WT and mutant CHIKV-SGR-D-Luc, ICRES-CHIKV, ONNV-2SG-ZsGreen, or WT CHIKV-SGR-nsP3-mCh-FF-Luc and WT CHIKV-SGR-FF-Luc were synthesized and purified using the mMACHINE™ SP6 Transcription Kit (Thermo Fisher Scientific). In brief, 250 ng of each of the RNAs was transfected into 5 x 10^4^ cells using Lipofectamine 2000 (Life Technologies) according to the manufacturer’s instructions. Cells were harvested and both FF-Luc and R-Luc activities were measured using the Dual-luciferase Kit (Promega) according to the manufacturer’s instructions. All experiments were repeated three times independently.

### Sequence analysis

Total RNA from transfected or infected cells was extracted with Trizol, followed by reverse transcription to generate cDNA using the SuperScript IV Reverse Transcriptase (Invitrogen) according to the manufacturer’s instructions. The expected DNA fragments of the CHIKV or ONNV genome were amplified by PCR (primer sequences available upon request) and analysed by Sanger sequencing.

### MTT assay

C2C12 cells were seeded into 96 well plates. The next day, cells were mock transfected and incubated at 37°C or 28°C. At indicated time points, medium was removed and replaced with 150 µl phenol-red free DMEM and 30 µl 6 mM MTT. Cells were incubated at 37°C for 2 h before media was removed and replaced with 100 µl DMSO. Plates were covered with foil and agitated until all MTT granules had dissolved, prior to reading the absorbance at 570 nm.

### ICRES-CHIKV one-step growth curve

C2C12 cells were seeded into 12 well plates. The next day, cells were infected with ICRES-CHIKV at MOI 0.1 pfu/cell and incubated at 37°C for 1 h. The cells were then washed with PBS and replaced with fresh complete media, followed by incubation at 37°C or 28°C respectively. At indicated time points cell supernatants were harvested for plaque assay, and cell were lysed and extracted with TRIzol solution (ThermoFisher) for qRT-PCR analysis.

### Plaque assay

Cell supernatants were stored at -80°C until required, thawed, serially diluted and applied to monolayers of BHK-21 cells for 1 h at 37°C. The inocula were aspirated and cells were overlaid with 0.8% methylcellulose for 72 h at 37°C or 96 h at 28°C for ONNV infection, and 48 h at 37°C for ICRES-CHIKV. Plaques were either visualized by EVOS microscopy (ONNV) or stained with 0.05% crystal violet for 30 min (CHIKV).

### qRT-PCR assay

Total RNA from transfected or infected cells was extracted with Trizol. Strand-specific CHIKV RNA primers were chosen from previous work [64] to amplify nucleotides 1-1560 of the CHIKV genome and used in reverse transcription reactions with a LunaScript8482 RT SuperMix Kit (NEB), followed by qPCR using the 2x qPCRBIO SyGreen Blue Mix (PCRBIOSYSTEMS). Positive and negative standards were generated using capped ICRES-CHIKV RNA with strand-specific RNA primers and were *in vitro* transcribed as described before.

### Western blotting

Cells were lysed in PLB or Glasgow Lysis Buffer (GLB) (10mM PIPES-NaOH pH7.2, 120mM KCl, 30mM NaCl, 5mM MgCl_2_, 1% Triton X-100, 10% glycerol) supplemented with protease inhibitors (Roche Diagnostics), protein samples were quantified by BCA assay (Pierce). 10 or 20 µg of each sample was heat denatured at 95°C for 5 min and separated by SDS-PAGE, proteins were then transferred to polyvinylidene fluoride (PVDF) membrane and blocked with 50% (v/v) Odyssey blocking buffer (Li-Cor) diluted in 1X Tris-buffered saline Tween-20 (TBS-T) (50 mM Tris-HCl pH 7.4, 150 mM NaCl, 0.1% Tween-20). The membrane was then incubated with primary antibodies (1:1000 dilution) at 4°C overnight and the next day incubated with fluorescently labelled anti-mouse (700 nm) and anti-rabbit (800 nm) secondary antibodies for 1 h at room temperature (RT). Membranes were then imaged on a Li-Cor Odyssey Sa Imager (Li-cor).

### Immunofluorescence assay

C2C12 cells were seeded into 12 well plates with coverslips. The next day, cells were transfected with WT CHIKV-SGR-nsP3-mCh-FF-Luc as described before. At indicated time points cells were fixed with 4% formaldehyde, washed with PBS, permeabilized with ice-cold methanol at -20 °C for 10 min and blocked with 2% BSA for 1 h. The cells were incubated with primary antibody to G3BP1 (Abcam) (1:200) at 4°C overnight. Cells were washed with DEPC-PBS, prior to incubation with J2 dsRNA antibody (Scicons) (1:100 dilution) at RT for 1 h. All cells were stained with Alexa Fluor-594 chicken anti-rabbit and Alexa Fluor-647 goat anti-mouse secondary antibodies (1:500 dilution) at RT for 1 h. Cells were mounted onto glass slides with Prolong Gold antifade reagent (Invitrogen) containing 4’,6’-diamidino-2-phenylindole dihydrochloride (DAPI) and sealed with nail varnish. Confocal microscopy images were acquired using a Zeiss LSM880 upright microscope. Post-acquisition analysis of images was performed using Zen software (Zen version 2018 black edition 2.3, Zeiss), or Fiji ImageJ (v1.49) software.

### Bulk RNA sequencing (RNA-Seq)

Capped RNAs of WT and GAA CHIKV-SGR-D-Luc were synthesized and purified as described above. In brief, 250 ng of each of the RNAs was transfected into 5 x 10^4^ cells using Lipofectamine 2000 (Life Technologies) according to the manufacturer’s instructions. Cells were harvested at 12 or 24 h.p.t. Starting from 50 ng total RNA per sample, bulk RNA barcoding and sequencing (BRB-Seq) was performed as described [65] with minor modifications. Briefly, reverse transcription was performed with Template Switching RT Enzyme Mix (NEB) and a uniquely barcoded oligo(dT)30 primer for each sample, modified to use Illumina TruSeq Read 1 priming site instead of Nextera Read 1 [66]. Following the remainder of the standard BRB-Seq protocol, up to 24 barcoded first-strand cDNAs were pooled into a single tube, second-strand cDNA was synthesized with Gubler-Hoffman nick translation, and tagmented cDNA pools with Tn5 produced in-house [67]. Fourteen cycles of PCR using a P5-containing primer and a unique multiplexed i7 indexing primer (Chromium i7 Multiplex Kit, 10X Genomics) were performed. Libraries were purified and size-selected with 0.65X SPRIselect (Beckman Coulter) and then 0.7X SPRIselect. All pools were sequenced on a NovaSeq SP v1.5 flowcell (Illumina) with 28 cycles of Read 1, 8 cycles of Read i7, and 101 cycles of Read 2.

### RNA-Seq analysis

Reads were demultiplexed with Picard v2.25.6 (from within viral-core v2.1.33) using the read structure “5S8B15M8B101T” and Q20M1 mismatch tolerance. Reads were mapped to the mouse genome mm39 with STAR v2.7.10a, and those reads mapping to genes on the primary assembly were counted with htseq-count v1.99.2. Pairwise differential gene expression between conditions was performed using DESeq2 (1.40.2) [68]. Genes whose expression changed 2-fold between conditions with adjusted *p*-value <0.05 (Wald test, Benjamini-Hochberg procedure for multiple hypothesis testing) were considered differentially expressed. The ISGs (dataset from HALLMARK_ALPHA_RESPONSE [50]) from Mus musculus species that were significantly expressed were labeled as triangles while the others were as dots. Heatmaps of the differential expressed ISGs at 37°C or 28°C were generated using the R package (R-studio v4.3.1).

### Data and code availability

Raw RNA-Seq reads and the processed counts matrix can be accessed at NCBI GEO accession: GSE264494. Reviewers may access this submission using the token ‘epglissetbkldmj’, which will be made public following publication. The analysis scripts used in this study are available at: [#https://github.com/path/to/repo#].

### Gene set enrichment analysis (GSEA)

GSEA was performed by using GSEA software version (4.3.2), downloaded from Broad Institute Gene Set Enrichment Analysis website (www.broad.mit.edu/gsea). The enrichment gene sets used in the analysis were selected in MSigDB (HALLMARK_ALPHA_RESPONSE) from Mus musculus species. Significance of enrichment magnitude was set at a False Discovery Rate (FDR) of 5% for GSEA.

### Statistical analysis

Statistical analysis was performed using the unpaired Student’s t tests (two-tailed). Data in bar graphs represent statistically significant differences and displayed as the means ± S.E. of three experimental replicates (n = 3).

## Acknowledgments

We thank the following people for their invaluable assistance in this work: Prof. Andres Merits (Institute of Technology, University of Tartu, Estonia) for the CHIKV SGRs, ICRES constructs, ONNV-2SG-ZsGreen construct and antibody to nsP1. Dr. Andrew Tuplin for the Zika virus replicon, Dr John Barr for the BUNV mini-genome construct, Drs. Niluka Goonawardane and Ruth Hughes for help in Fiji ImageJ analysis, Dr Sreenivasan Ponnambalam for access to the EVOS microscope (all at the University of Leeds). Dr. Wei Wang, Jennifer Miller, and Jean Arly Volmar of the Genomics Core Facility at the Princeton University Lewis-Sigler Institute for Integrative Genomics for next-generation sequencing assistance, Drs. Abhishek Biswas and Lance R. Parsons for the help with the RNA-seq analysis.

