## Supplementary figures S1-S7 for "Enhancement of Chikungunya virus genome replication in mammalian cells at a sub-physiological temperature"

### Slide 1
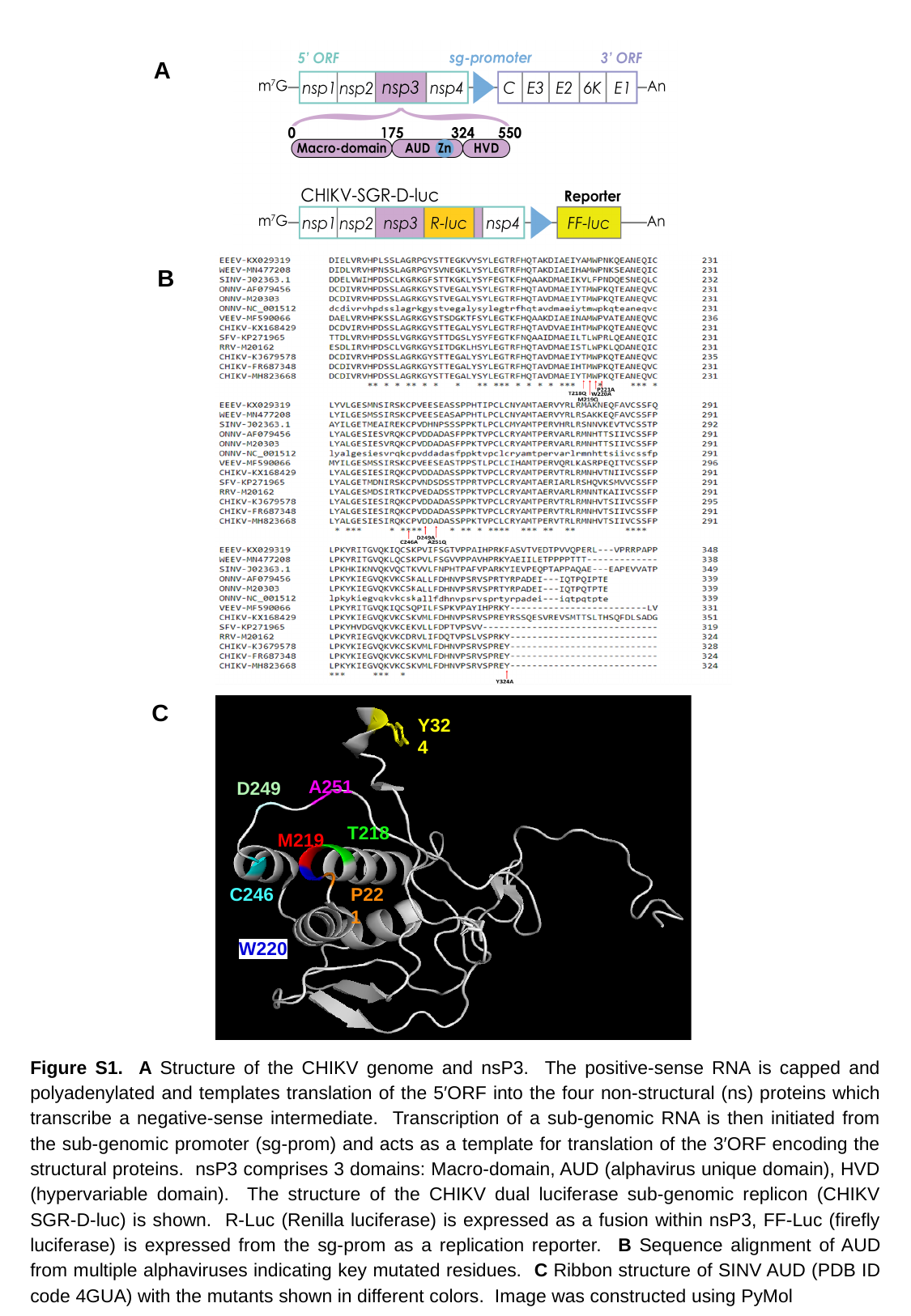

A
B
C
Y324
A251
D249
T218
M219
P221
C246
W220
Figure S1. A Structure of the CHIKV genome and nsP3. The positive-sense RNA is capped and polyadenylated and templates translation of the 5′ORF into the four non-structural (ns) proteins which transcribe a negative-sense intermediate. Transcription of a sub-genomic RNA is then initiated from the sub-genomic promoter (sg-prom) and acts as a template for translation of the 3′ORF encoding the structural proteins. nsP3 comprises 3 domains: Macro-domain, AUD (alphavirus unique domain), HVD (hypervariable domain). The structure of the CHIKV dual luciferase sub-genomic replicon (CHIKV SGR-D-luc) is shown. R-Luc (Renilla luciferase) is expressed as a fusion within nsP3, FF-Luc (firefly luciferase) is expressed from the sg-prom as a replication reporter. B Sequence alignment of AUD from multiple alphaviruses indicating key mutated residues. C Ribbon structure of SINV AUD (PDB ID code 4GUA) with the mutants shown in different colors. Image was constructed using PyMol

### Slide 2
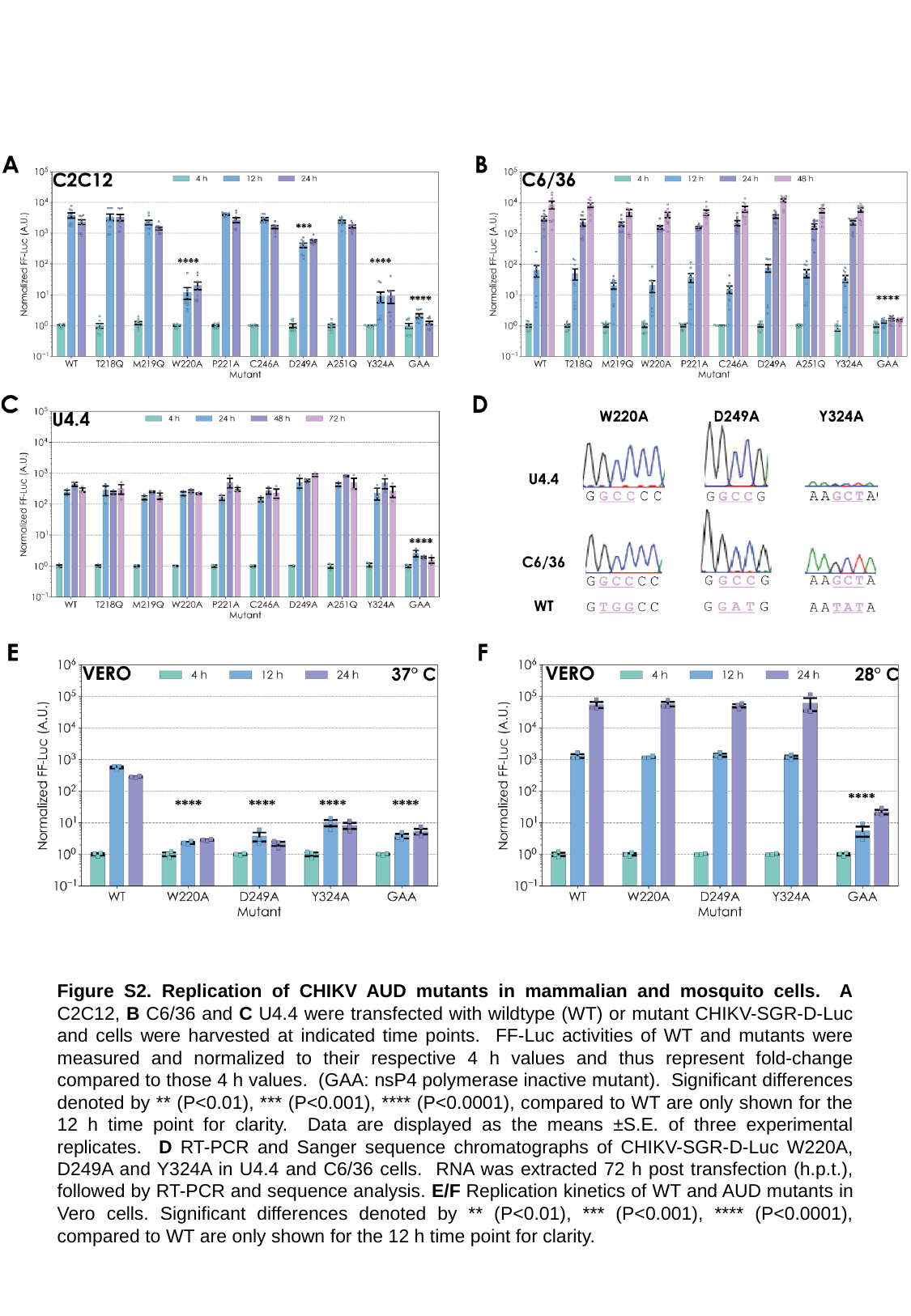

Figure S2. Replication of CHIKV AUD mutants in mammalian and mosquito cells. A C2C12, B C6/36 and C U4.4 were transfected with wildtype (WT) or mutant CHIKV-SGR-D-Luc and cells were harvested at indicated time points. FF-Luc activities of WT and mutants were measured and normalized to their respective 4 h values and thus represent fold-change compared to those 4 h values. (GAA: nsP4 polymerase inactive mutant). Significant differences denoted by ** (P<0.01), *** (P<0.001), **** (P<0.0001), compared to WT are only shown for the 12 h time point for clarity. Data are displayed as the means ±S.E. of three experimental replicates. D RT-PCR and Sanger sequence chromatographs of CHIKV-SGR-D-Luc W220A, D249A and Y324A in U4.4 and C6/36 cells. RNA was extracted 72 h post transfection (h.p.t.), followed by RT-PCR and sequence analysis. E/F Replication kinetics of WT and AUD mutants in Vero cells. Significant differences denoted by ** (P<0.01), *** (P<0.001), **** (P<0.0001), compared to WT are only shown for the 12 h time point for clarity.

### Slide 3
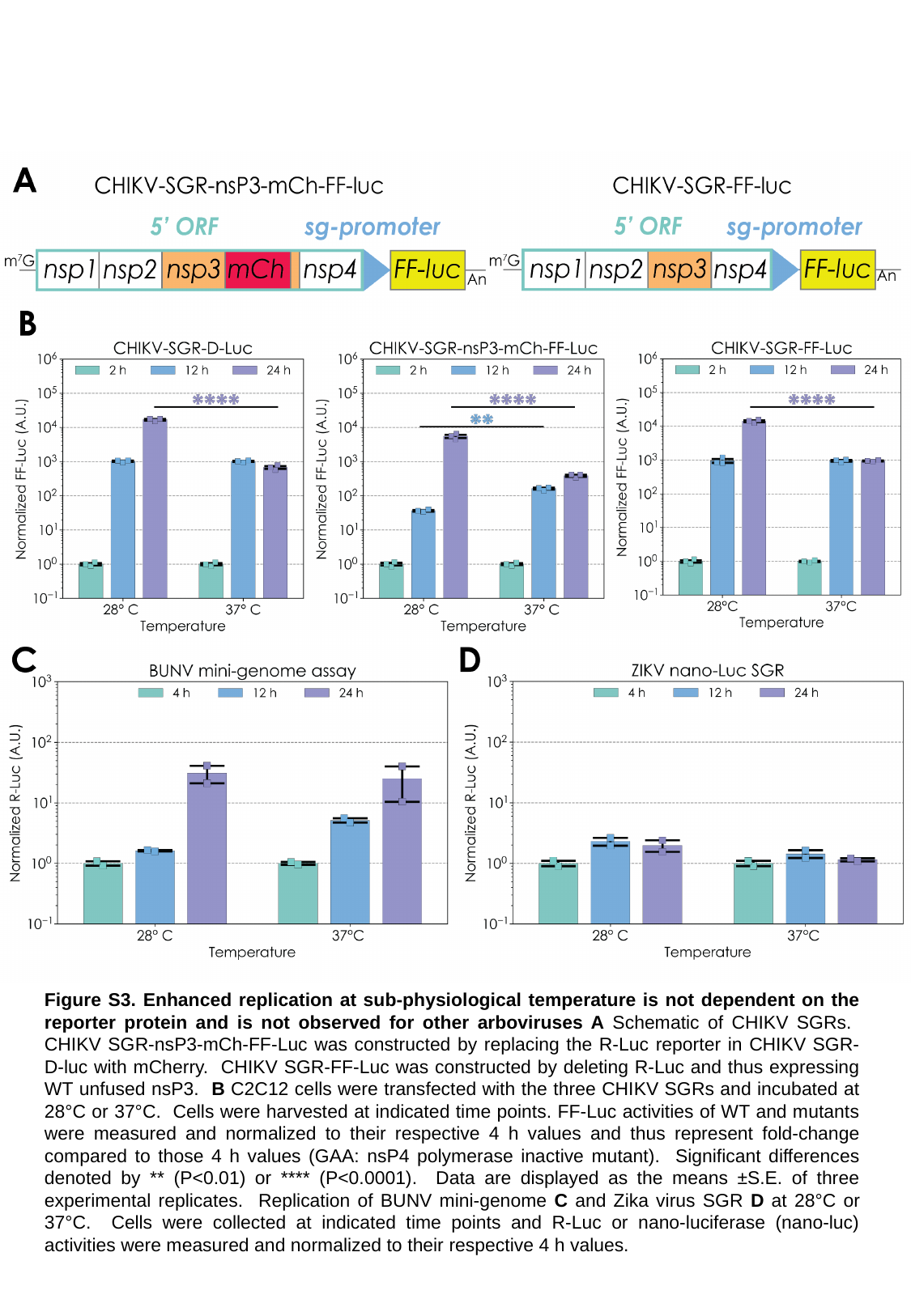

Figure S3. Enhanced replication at sub-physiological temperature is not dependent on the reporter protein and is not observed for other arboviruses A Schematic of CHIKV SGRs. CHIKV SGR-nsP3-mCh-FF-Luc was constructed by replacing the R-Luc reporter in CHIKV SGR-D-luc with mCherry. CHIKV SGR-FF-Luc was constructed by deleting R-Luc and thus expressing WT unfused nsP3. B C2C12 cells were transfected with the three CHIKV SGRs and incubated at 28°C or 37°C. Cells were harvested at indicated time points. FF-Luc activities of WT and mutants were measured and normalized to their respective 4 h values and thus represent fold-change compared to those 4 h values (GAA: nsP4 polymerase inactive mutant). Significant differences denoted by ** (P<0.01) or **** (P<0.0001). Data are displayed as the means ±S.E. of three experimental replicates. Replication of BUNV mini-genome C and Zika virus SGR D at 28°C or 37°C. Cells were collected at indicated time points and R-Luc or nano-luciferase (nano-luc) activities were measured and normalized to their respective 4 h values.

### Slide 4
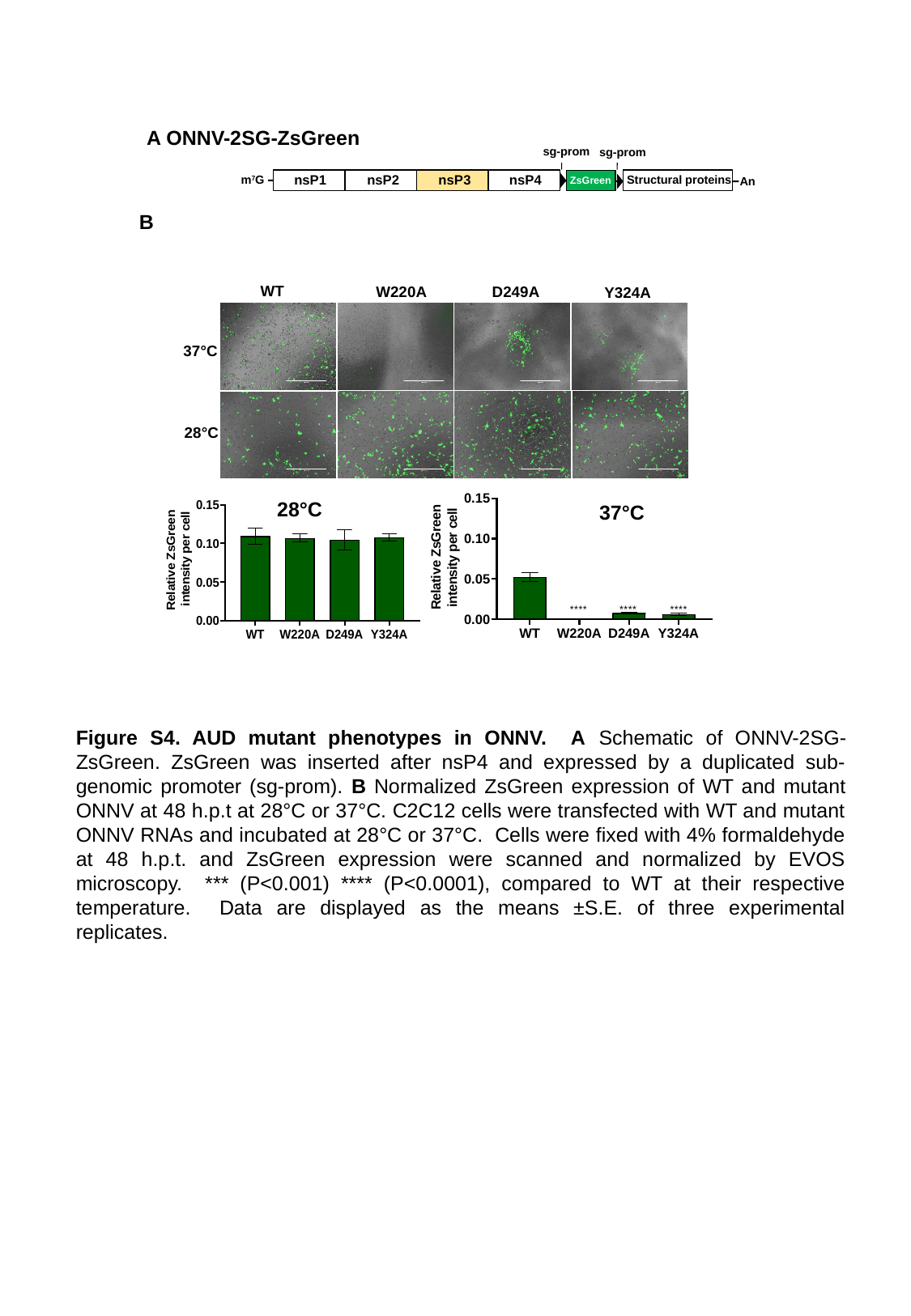

A ONNV-2SG-ZsGreen
sg-prom
sg-prom
nsP1
nsP2
nsP3
nsP4
m^7^G
Structural proteins
ZsGreen
An
B
WT
W220A
D249A
Y324A
37°C
28°C
28°C
37°C
Figure S4. AUD mutant phenotypes in ONNV. A Schematic of ONNV-2SG-ZsGreen. ZsGreen was inserted after nsP4 and expressed by a duplicated sub-genomic promoter (sg-prom). B Normalized ZsGreen expression of WT and mutant ONNV at 48 h.p.t at 28°C or 37°C. C2C12 cells were transfected with WT and mutant ONNV RNAs and incubated at 28°C or 37°C. Cells were fixed with 4% formaldehyde at 48 h.p.t. and ZsGreen expression were scanned and normalized by EVOS microscopy. *** (P<0.001) **** (P<0.0001), compared to WT at their respective temperature. Data are displayed as the means ±S.E. of three experimental replicates.

### Slide 5
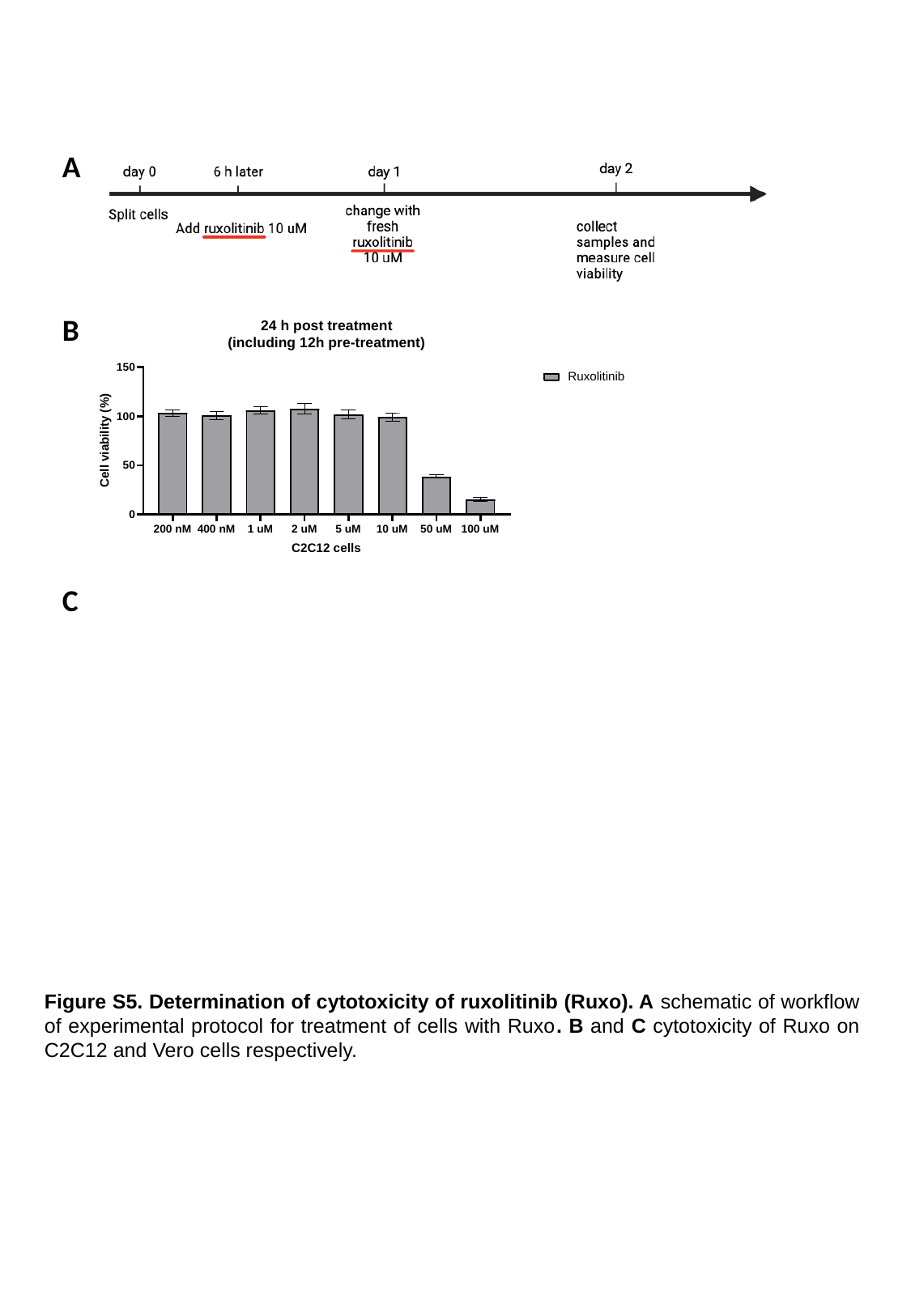

A
B
C
Figure S5. Determination of cytotoxicity of ruxolitinib (Ruxo). A schematic of workflow of experimental protocol for treatment of cells with Ruxo. B and C cytotoxicity of Ruxo on C2C12 and Vero cells respectively.

### Slide 6
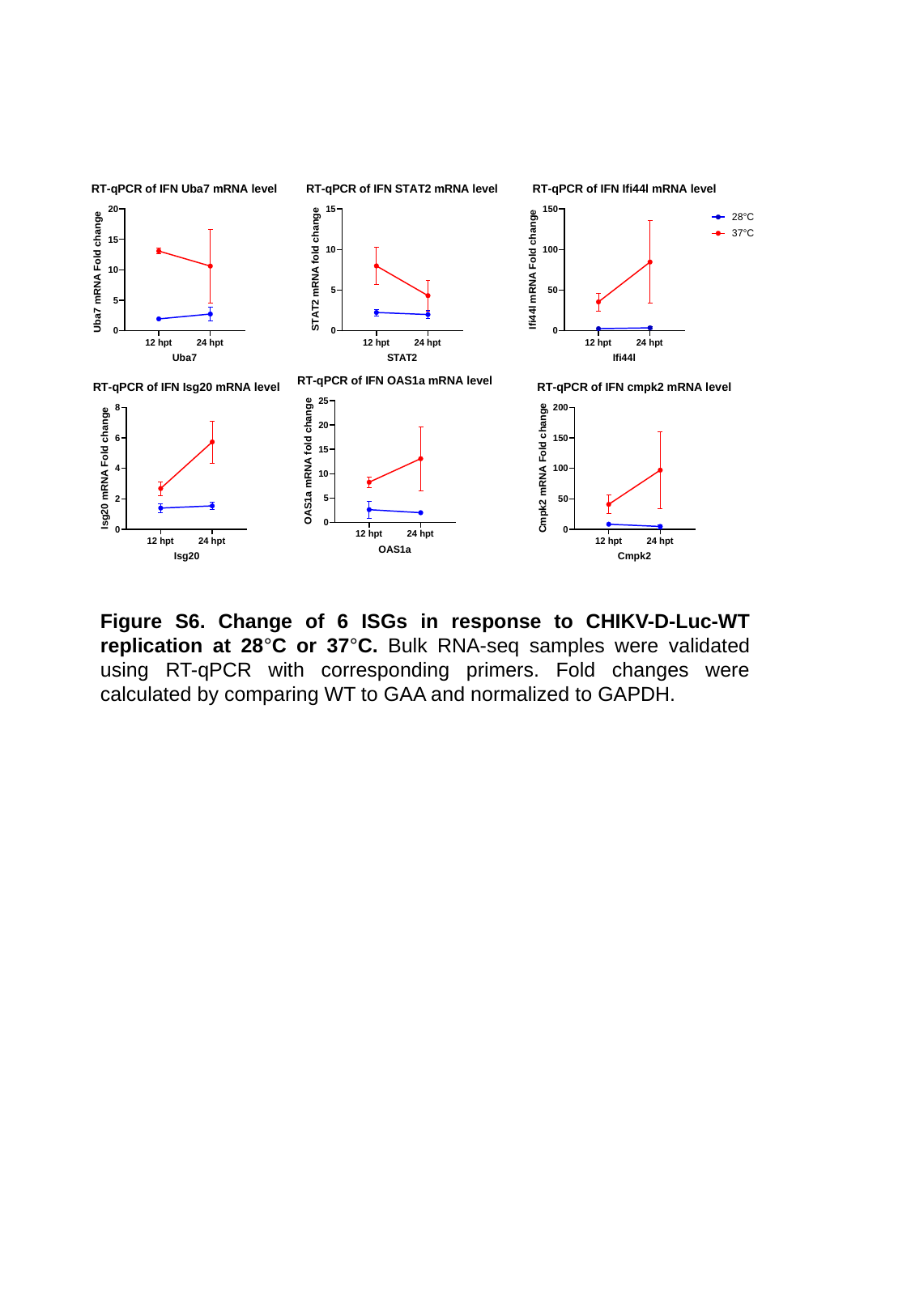

Figure S6. Change of 6 ISGs in response to CHIKV-D-Luc-WT replication at 28°C or 37°C. Bulk RNA-seq samples were validated using RT-qPCR with corresponding primers. Fold changes were calculated by comparing WT to GAA and normalized to GAPDH.

### Slide 7
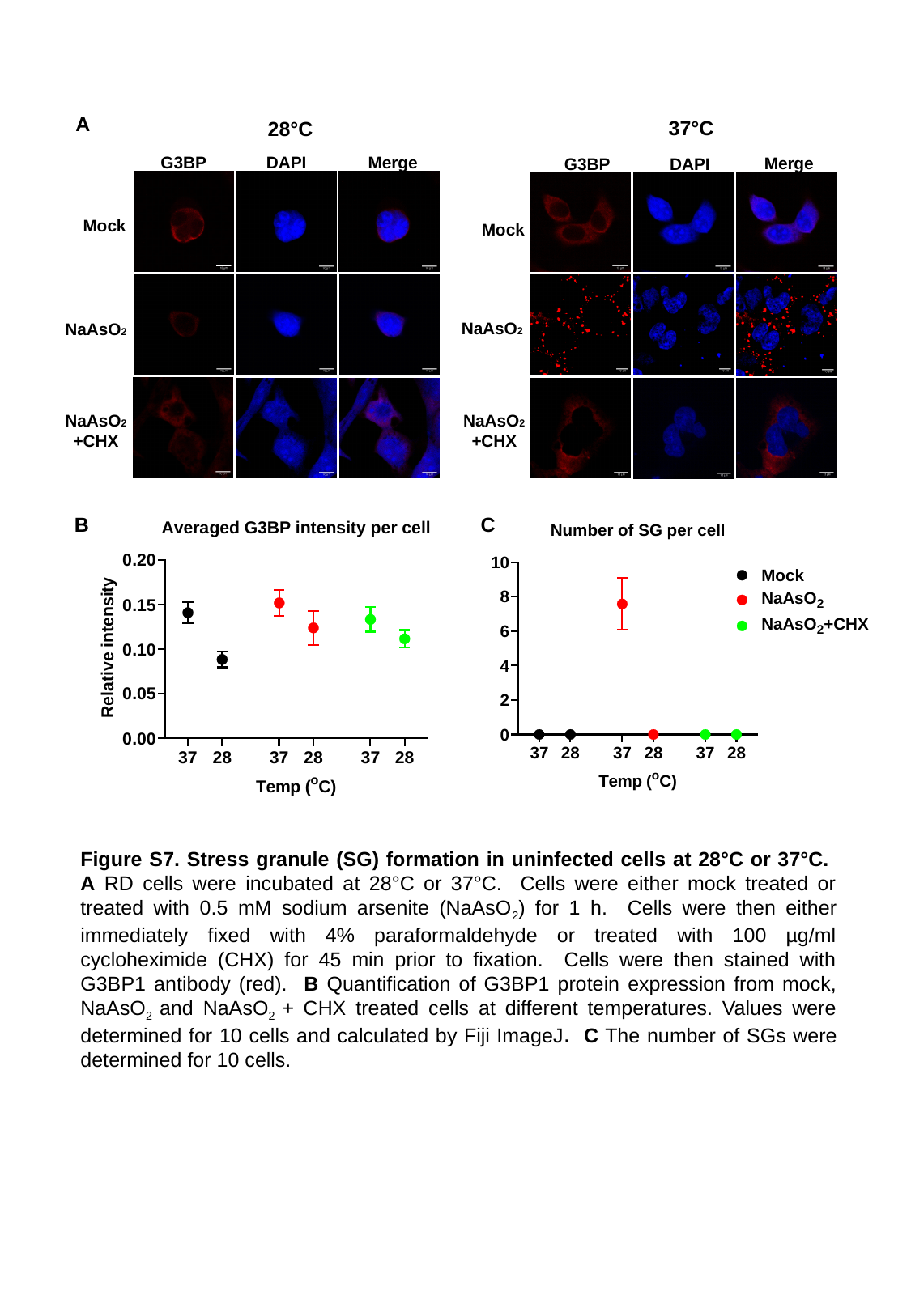

A
37°C
Merge
G3BP
DAPI
Mock
NaAsO2
28°C
G3BP
DAPI
Merge
Mock
NaAsO2
NaAsO2
+CHX
NaAsO2
+CHX
B
C
Figure S7. Stress granule (SG) formation in uninfected cells at 28°C or 37°C. A RD cells were incubated at 28°C or 37°C. Cells were either mock treated or treated with 0.5 mM sodium arsenite (NaAsO_2_) for 1 h. Cells were then either immediately fixed with 4% paraformaldehyde or treated with 100 µg/ml cycloheximide (CHX) for 45 min prior to fixation. Cells were then stained with G3BP1 antibody (red). B Quantification of G3BP1 protein expression from mock, NaAsO_2_ and NaAsO_2_ + CHX treated cells at different temperatures. Values were determined for 10 cells and calculated by Fiji ImageJ. C The number of SGs were determined for 10 cells.
